# Identification of cyclic-di-GMP-modulating protein residues by bi-directionally evolving a social trait in *Pseudomonas fluorescens*

**DOI:** 10.1101/2022.05.10.491292

**Authors:** Collin Kessler, Wook Kim

## Abstract

Modulation of the intracellular cyclic di-guanosine monophosphate (c-di-GMP) pool is central to the formation of structured bacterial communities. Genome annotations predict the presence of dozens of conserved c-di-GMP catalytic enzymes in many bacterial species, but the functionality and regulatory control of the vast majority remain underexplored. Here, we begin to fill this gap by utilizing an experimental evolution system in *Pseudomonas fluorescens* Pf0-1, which repeatedly produces a unique social trait through bidirectional transitions between two distinct phenotypes converging on c-di-GMP modulation. Parallel evolution of 33 lineages captured 147 unique mutations among 191 evolved isolates in genes that are empirically demonstrated, bioinformatically predicted, or previously unknown to impact the intracellular pool of c-di-GMP. Quantitative chemistry confirmed that each mutation causing the phenotypic shift predictably amplifies or reduces c-di-GMP production. We integrate our mutation, phenotype, and quantification data with current models of known regulatory and catalytic systems, describe a previously unknown relationship between a regulatory component of branched-chain amino acid biosynthesis and c-di-GMP production, and predict functions of unexpected proteins that clearly impact c-di-GMP production. Sequential mutations that continuously disrupt or recover c-di-GMP production across discrete functional elements suggest a complex and underappreciated interconnectivity within the c-di-GMP regulome of *P. fluorescens*.

**Importance:** Microbial communities comprise densely packed cells where competition for space and resources is fierce. In our model system, mutant cells with a dry (D) phenotype are selected from a population with a mucoid (M) phenotype, and vice versa, because M and D cells physically work together to spread away from the overcrowded colony. D cells produce high levels of c-di-GMP and M cells produce low levels, so each mutation impacts c-di-GMP production. C-di-GMP is a second messenger which regulates diverse bacterial phenotypes that cause tremendous clinical and environmental problems. Many bacteria possess dozens of enzymes that are predicted to produce c-di-GMP, but most are considered to be non-functional. Here, we take advantage of the bi-directional selection of M and D phenotypes to identify key residues that could force these enzymes to turn on or off. Several unexpected proteins also participate in this process, but very little is known about them.

## Introduction

Microbes utilize diverse signaling mechanisms to adapt and respond to a dynamic environment. Cyclic di-guanosine monophosphate (c-di-GMP) is a ubiquitous secondary messenger in bacteria and high intracellular levels of c-di-GMP characteristically repress flagellar motility while stimulating the expression of biofilm-formation genes (1-4). Numerous c-di-GMP binding proteins and riboswitches respond to intracellular levels of c-di-GMP to also regulate cellular morphology, virulence factors, antibiotic production, and the cell cycle (5, 6). C-di-GMP is synthesized from two molecules of GTP by a diguanylate cyclase (DGC) and hydrolyzed by a phosphodiesterase (PDE) (1). DGCs and PDEs are identified by the GGDEF or EAL/HD-GYP domains, respectively, and there are also hybrid proteins with both DGC and PDE functions (7, 8). *Pseudomonas* spp. typically possess greater than 50 predicted DGCs or PDEs (9). Specific signaling pathways or regulators have been characterized to independently regulate several DGCs and PDEs (10-17), and some have been demonstrated to be under allosteric control, where a certain range of c-di-GMP levels either initiate or prevent enzymatic activity (18-22). However, the function of the vast majority of bioinformatically predicted DGCs and PDEs have not been empirically validated and their regulatory mechanisms remain unknown (23).

A previous experimental evolution study described the bidirectional evolution of mucoid (M) and dry (D) colony phenotypes in *P. fluorescens* Pf0-1, which likely occurs through c-di-GMP oscillation (24). An aging colony of M cells repeatedly produces spreading fans at the perimeter, which comprise the original M cells along with new D mutant cells that produce a dry and wrinkly colony morphology on their own. Additionally, an aging colony of D cells also repeatedly generates spreading fans, which comprise the original D cells along with a new mutant that appears phenotypically identical to the ancestral M. This bidirectional selection of M and D phenotypes occurs continuously because M and D cells self-organize in space and physically work together to spread out from the nutrient-poor and crowded conditions of dense colony growth, and neither type is capable of spreading out on their own. Genetic analyses of five evolved isolates revealed characteristic mutations in *wsp* genes to suggest that c-di-GMP production is likely elevated in D and reduced in M (24).

Here, we carry out a large-scale parallel evolution experiment using the bidirectional M-D system to identify 147 unique genes that are empirically demonstrated, bioinformatically predicted, or previously unknown to impact c-di-GMP levels. We expand the current model of c-di-GMP production by the Wsp signal transduction system and identify unique missense mutations outside the conserved catalytic domain of numerous DGCs that stimulate enzymatic activity. We also describe proteins with no previous associations to c-di-GMP production that appear to influence the activity of several DGCs, including a regulatory component of the branched-chain amino acid (BCAA) biosynthesis pathway. Lastly, we map the interconnectivity of the c-di-GMP regulome through sequential mutations that disrupt or recover c-di-GMP production across functionally discrete systems.

## Results and Discussion

### Bidirectional evolution of mucoid and dry phenotypes occurs through intracellular c-di-GMP pool oscillation

The bidirectional selection of mucoid (M) and dry (D) phenotypes occurs continuously in our experimental system (24) because neither type is capable of spreading out on their own, but a colony of initially mixed M and D cells readily spreads out in a radial pattern without requiring additional mutations (Fig. 1A). With the expectation that a large-scale utilization of this experimental evolution system could identify diverse mechanisms of c-di-GMP modulation, we initially set up 25 parallelly evolving lineages of M. As anticipated, we observed bidirectional transitions between M and D phenotypes across all lineages. In total, we isolated over 600 strains, analyzed genome sequences of 191 strains, and quantified c-di-GMP in a subset of 56 strains. Remarkably, we detected a single mutation associated with each phenotypic transition, with the exception of only few additional silent mutations across all genome sequences (Fig. S1, Table S1, Table S1 – Source Data 1). Genome sequencing successive strains within each lineage provided an internal control, which confirmed the exclusive presence of the respective preceding mutations.

**Figure 1.**
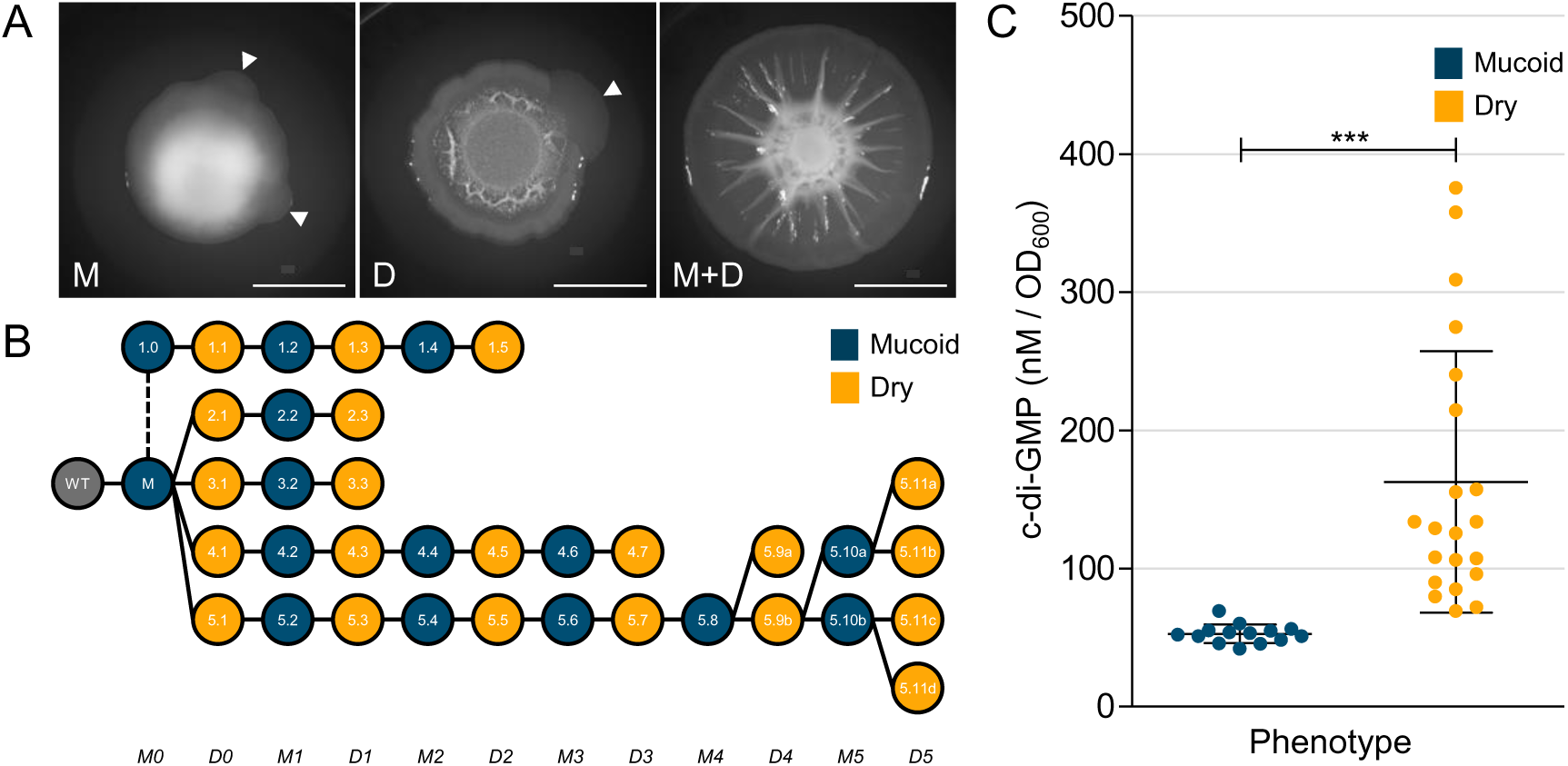
Bidirectional evolution of M and D phenotypes are c-di-GMP dependent. A) Colony morphologies of the ancestral mucoid (M, left) and dry (D, middle) strains after 4 days of growth. Arrows indicate the emergence of spreading fans that contain the respective parent strain and a new mutant strain with the opposite phenotype. The spreading phenotype is readily produced by artificially mixing M and D strains (right, scale bar = 10mm). B) Phylogeny of strains in select lineages colored by phenotype and corresponding colony morphologies are shown in Fig. S2. The numbers indicate the IDs of the evolved strains as shown in Table S1. Strain 1.0 was constructed from the ancestral M strain to lack the *wspC* gene, which encodes a methyltransferase known to activate the Wsp system. Each strain was isolated from naturally emerging fans from their respective parent colony with the opposite phenotype. This phylogeny depicts the bidirectional transitions between mucoid and dry states which we show are c-di-GMP dependent. C) C-di-GMP levels of evolved strains. C-di-GMP was quantified through LC-MS/MS and is reported as nM per OD600. Samples are categorized by phenotype and we report the triplicate mean ± SD for each phenotypic classification: mucoid n=14 and dry n=21 (*** Student’s t-test p < 0.001).

**Table 1.**
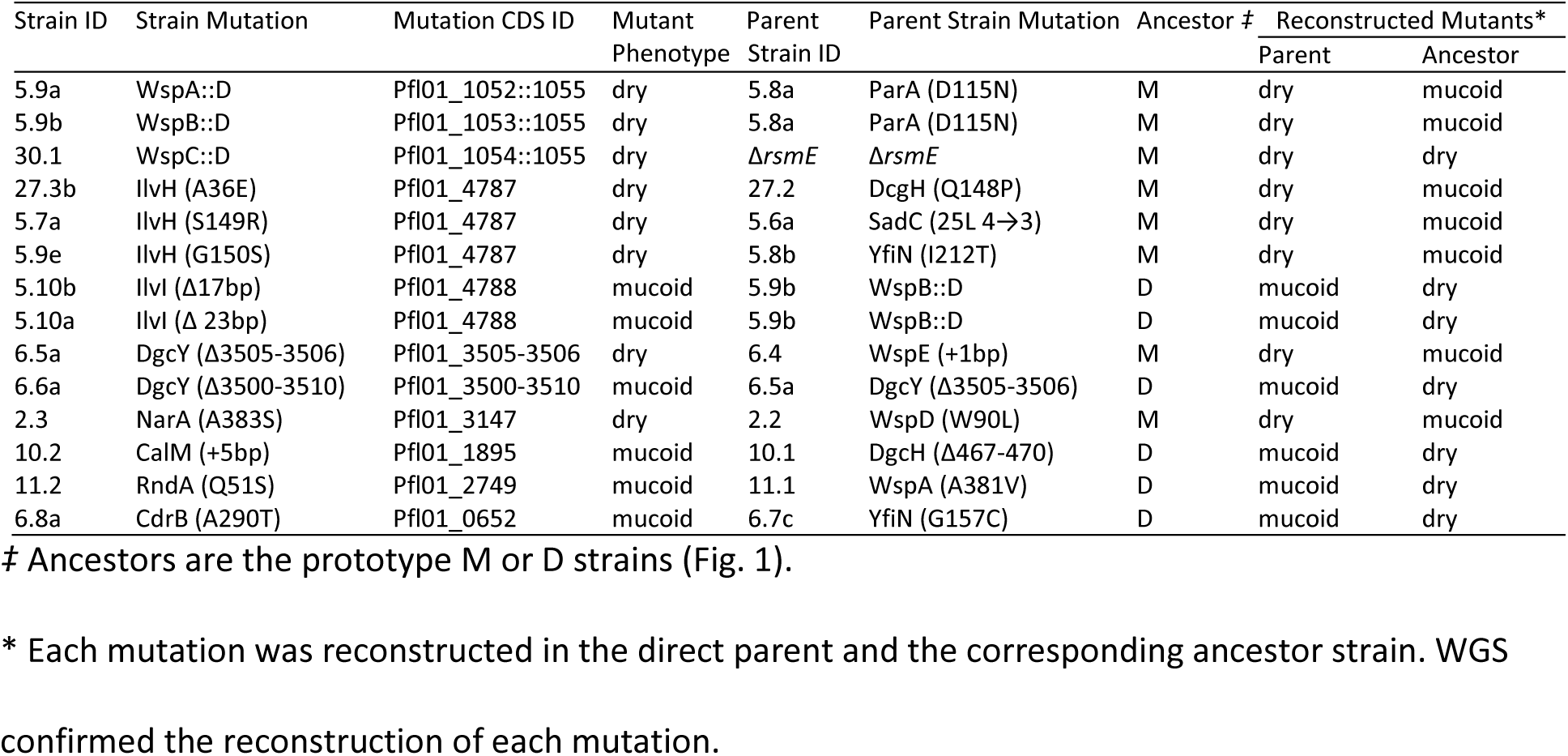
Reconstructing select naturally acquired mutations in both the ancestral and direct parent strains reveal dependence on the previous mutation(s) to drive specific M-D phenotypic shifts.

Among the 147 unique mutations detected, 81 mutations were in *wsp* genes and 66 mutations were in other genes including those that are hypothetical. Motility assays showed that all D strains were non-motile and all M strains were motile as expected, and LC-MS/MS quantification confirmed that c-di-GMP was elevated in all D strains relative to their respective parent M strain (Table S1). Notably, many mutations in non-*wsp* genes sequentially flanked specific mutations in the Wsp signal transduction system, the most extensively studied c-di-GMP regulatory system in *Pseudomonas* spp. (17). Several missense mutations observed in Wsp proteins identified in this study overlap with comparable mutations from various *Pseudomonas* and *Burkholderia* studies that we had previously compiled from the literature (17) (Fig. 2). However, the vast majority of our mutations had never been observed and many occur outside functionally annotated domains. The data presented in Fig. 2 represent the most comprehensive record of functionally important residues of the Wsp system to date and empower our predictions of mutation consequences. We first focus on sequential mutations in the Wsp system to reveal both predictable and surprising evolutionary innovations to modulate c-di-GMP production. For the sake of an accessible narrative, we will primarily refer to five simplified lineages (Fig. 1B), with corresponding c-di-GMP measurements (Fig. 1C) and colony morphologies (Fig. S2).

**Figure 2.**
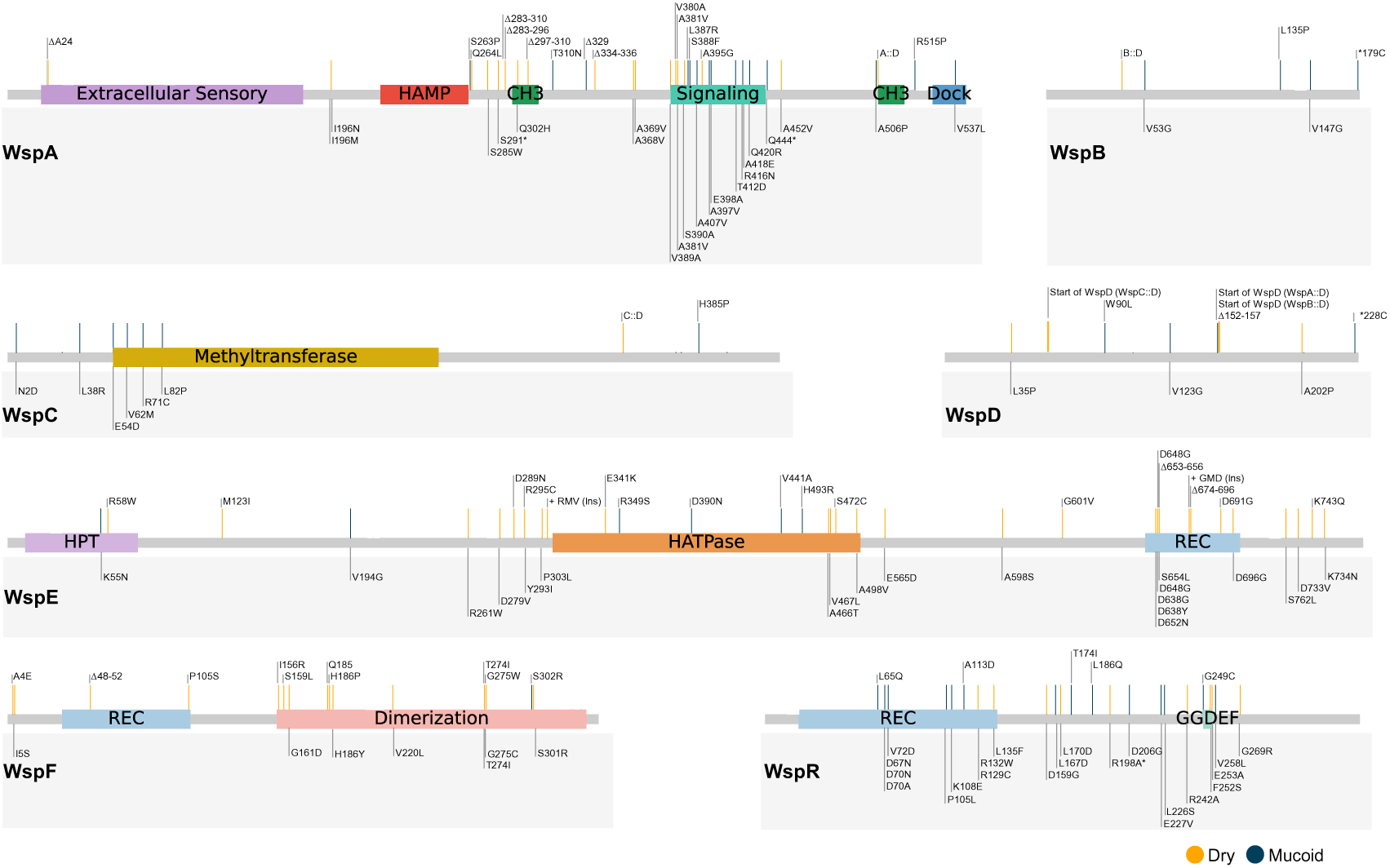
Alignment of our missense and in-frame deletion mutation data with previously annotated mutations in the Wsp system. Individual Wsp proteins are shown with their functional domains that were derived from CDD (NCBI), Prosite, or homology to the *E. coli* chemotaxis system (HAMP = transmembrane relay domain, CH3 = methylation site, HPT = histidine phospho-transfer domain, HATPase = histidine kinase ATP binding domain, and REC = phosphoreceiver domain). Only the missense and in-frame mutations identified in this study are shown above each protein and they are mapped to the consensus sequence of each Wsp protein (17). Previously annotated mutations (17) are shown within the gray box below each protein. Mutations indicated within the gray box are annotated to represent specific residues positions that were affected in the respective protein sequence of a given organism, but the grey horizontal bars indicate their actual positions within the consensus sequence. Therefore, the horizontal bar locations of the mutations identified in this study are directly comparable to previously identified mutations through the consensus sequence. Mutations that stimulate c-di-GMP production are indicated by orange horizontal bars and those that reduce c-di-GMP production are indicated by blue horizontal bars.

### Diverse mutations converge on the Wsp signal transduction system that dynamically rewire signal flow to produce a spectrum of functional states

The Wsp signal transduction system was first reported two decades ago in *P. fluorescens* (25) and continues to serve as the primary model system of c-di-GMP regulation in pseudomonads. The Wsp system comprises seven proteins encoded in a monocistronic operon under the control of an unidentified transcriptional regulator (26), and the current functional model is based on a combination of empirical studies and predictions based on homologies to the chemotaxis system in *Escherichia coli* (17). The main difference with the latter is that the signal cascade is initiated by the extracellular sensory domain of WspA detecting surface contact (27, 28) to ultimately drive c-di-GMP production by the DGC WspR (Fig. 3). Given that a mutation in any one of the seven Wsp protein coding genes could potentially increase or decrease c-di-GMP production through WspR, we expected *wsp* genes to be a common target in our parallel evolution experiments. Indeed, the vast majority of the first transitions from the ancestral M to D occurred through either a missense or an in-frame deletion mutation in *wsp* genes (Table S1).

**Figure 3.**
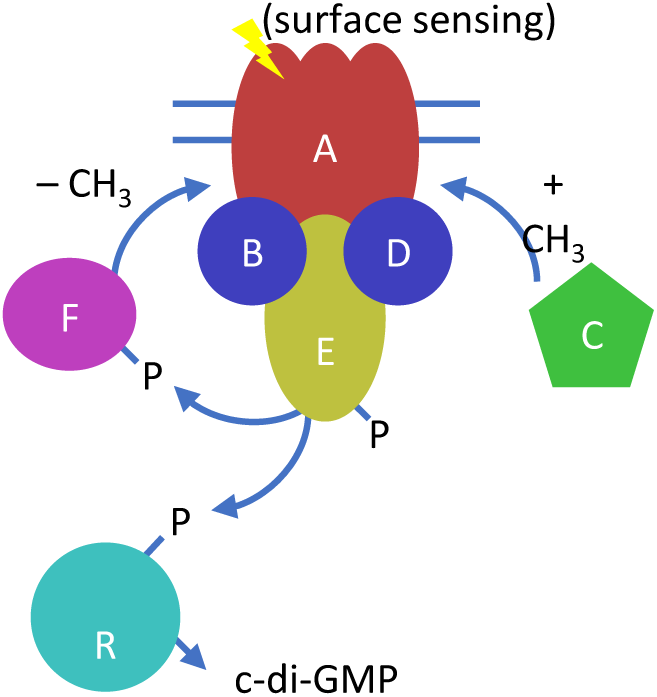
Functional model of the Wsp signal transduction system. The Wsp system in *P. aeruginosa* responds to surface contact to ultimately produce c-di-GMP. (CH3 = methyl group, P= phosphoryl group). Homology with the chemotaxis system predicts that once WspA is stimulated by surface-contact, it undergoes a conformational change to expose cytosolic methylation helices that are necessary for signal transduction (29). The methyl-transferase WspC is then predicted to dock at the 5’ tail of WspA and methylate the WspA trimer-of-dimers complex (56). The signal is then relayed through the coupling proteins WspB and WspD, and subsequently to WspE, which functions as a histidine kinase (57). WspE phosphorylates the DGC WspR to stimulate c-di-GMP production (58). Simultaneously, WspE phosphorylates the methyl-esterase WspF, which de-methylates WspA and terminates the signaling cascade (26). Significant work has been carried out to characterize the biochemical functions of WspE and WspR in particular, and WspE is a common mutational target in a clinical setting (17). More recently, studies have begun to explore the functional role of WspA during signal activation and subcellular localization, revealing that both properties are dependent on WspB and WspD proteins (27, 59). These studies also suggest that WspB and WspD likely possess unique functions, but they remain unresolved.

Across our subset of five lineages (Fig. 4), initial mutations appear to artificially activate WspA, followed by surprisingly similar patterns of sequential mutations in various *wsp* genes to oscillate c-di-GMP production. Lineages 2 and 3 began with the first respective D strain possessing a unique in-frame deletion that partially spans the predicted methylation helices of WspA. Lineage 5 began with a missense mutation in the phosphoreceiver domain of WspF (P105S), which likely disrupts its methyl-esterase activity to keep WspA in an activated state. This activation is undone in the following M strain by an in-frame deletion in *wspA* that removes the amino acid residue S329, then c-di-GMP production is restored again in the next D strain through an in-frame deletion of residues 282-296 in WspA. This deletion falls within the same deleted region as those observed in lineages 2 and 3 (Fig. 2, Fig. 4).

**Figure 4.**
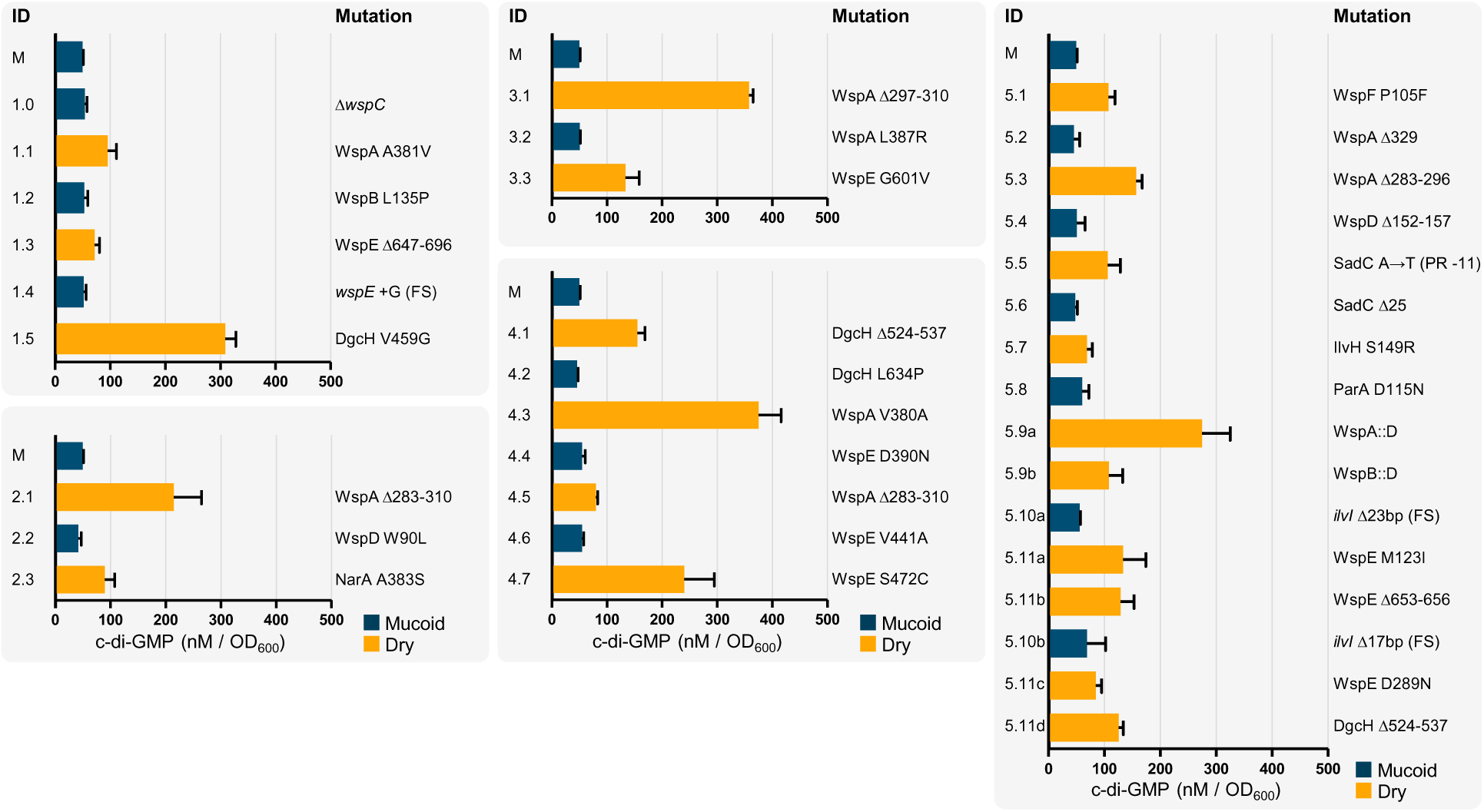
C-di-GMP oscillates between all M and D phenotypic transitions. C-di-GMP was quantified via LC-MS/MS and is reported as the triplicate mean ± SD. The charts capture the independently evolved strains within the five selected lineages shown in Figure 1B. D strains are shown in orange and M strains are shown in blue and their respective colony morphologies are shown in Fig. S2. The numerical IDs correspond to Table S1, and mutations reflect the affected amino acid residues, except frameshift (FS) and promoter region (PR) mutations refer to the specific nucleotides.

Although methylation of WspA remains to be empirically demonstrated, its predicted methylation helices clearly have a functional role in Wsp signaling as they were commonly mutated to activate c-di-GMP production (Fig. 2). Based on our mutation data and homologies to Tsr-methylation in the chemotaxis system of *E. coli* (29), we predict that the residues Q285, E292, E299, Q502, and E512 are methylated and the deletion of these sites mimics a methylated state. We also set up an independent lineage to commence with an engineered *ΔwspC* M strain (Fig. 4, strain 1.0), under the expectation that the deletion of *wspC* would steer downstream mutations away from the Wsp system. However, we detected a missense mutation within the signaling domain of WspA (A381V) in the first D progeny strain 1.1, and the same A381V mutation was previously observed in an independently evolved D strain with an unaltered *wspC* gene (24). We detected additional missense mutations in WspA’s signaling domain to independently stimulate c-di-GMP production (Fig. 2), which likely bypass the methylation requirement to activate WspE.

Mutations that stimulate WspA were typically followed by a missense mutation in WspB, WspD, or WspE (Table S1). A missense mutation in WspD (W90L) in lineage 2 reduced c-di-GMP production like the WspB (L136P) mutation in lineage 1 (Fig. 4). The functional roles of WspB and WspD in signal relay are loosely modeled after homology to CheW of *E. coli*, which remains as the least characterized protein in the chemotaxis system. Mutations in CheW that successfully disrupt signal transduction are largely limited to five unique amino acid clusters that situate on the protein surface and are essential for the docking of CheW to Tsr (WspA homolog) (30). WspB and WspD also possess six and seven highly conserved amino acid clusters, respectively (17). Our WspB L135P and WspD W90L mutations fall within these conserved clusters that likely situate on the surface of WspB and WspD and natively act as docking sites for WspA and WspE to permit signal relay. We also observed additional missense and in-frame deletion mutations in WspB and WspD that independently impact c-di-GMP production (Fig.2, Table S1), confirming that both WspB and WspD are necessary for Wsp signal transduction through domains and interactions that remain to be characterized.

Loss-of-function mutations (i.e. nonsense and frameshift mutations) in *wspE* were exclusively associated with D to M transitions throughout all lineages to reduce c-di-GMP production (Table S1). These observations are not particularly surprising, since WspR cannot be phosphorylated in the absence of a functional WspE (Fig. 3). In contrast, multiple missense and in-frame deletions were observed in the phosphoreceiver domain of WspE to exclusively stimulate c-di-GMP production (Fig. 2). Accordingly, mutations within the conserved phosphoreceiver domain of CheY, a WspE homolog in *Escherichia coli*, can activate the protein without phosphorylation (31, 32). In addition, the DGC activity of WspR in *P. aeruginosa* could be modulated in a phosphorylation-independent manner (16, 33). Artificial activation of WspE would deregulate WspA-dependent activation, and was often the evolutionary solution to recover c-di-GMP production when the upstream signaling components of the Wsp system became functionally crippled (Table S1). The M strain 1.4 of lineage 1 carries a frameshift mutation in *wspE*, which should terminally shut down the Wsp system, and the c-di-GMP level was reduced as predicted (Fig. 4). However, c-di-GMP production was restored in this lineage once again in the following D strain 1.5 through a missense mutation in DgcH (V459G), a bioinformatically predicted DGC that impacts biofilm formation in *P. aeruginosa*PAO1 (34). DgcH is not only capable of producing c-di-GMP, but at one of the highest levels detected in our study (Fig. 4, Table S1). In contrast to the Wsp system, it is difficult to predict functional consequences of mutations in other DGCs, since so little is known beyond their conserved catalytic domain.

### Many DGCs are functionally capable of causing the M to D shift independently from WspR

Sequential patterns of mutations and c-di-GMP quantification across all 33 lineages indicate that the DGC DgcH is also a dominant contributor to the intracellular c-di-GMP pool under our experimental conditions. Other known or predicted DGCs were typically targeted to modulate c-di-GMP levels after the Wsp system and DgcH became genetically dysfunctional. One exception is MorA, which represents the first mutation target in two lineages (Table S1). The first D strain 4.1 of lineage 4 carries an in-frame deletion mutation in *dgcH* that removes amino acid residues 524-537 (Fig. 4, Fig. 5). Interestingly, the GGDEF diguanylate cyclase domain of DgcH spans residues 537-693 and the removal of the immediately adjacent residues appears to activate c-di-GMP production. We observed a reduction in c-di-GMP in the following M strain 4.2, which has a missense mutation in DgcH (L634P). A search against NCBI’s conserved domain database identifies that the residues 619-643 contain the inhibitory I-site (22, 35), with residue N637 as the predicted interaction site with c-di-GMP. Our mutation data supports this annotation and the likelihood that DgcH is under allosteric control by c-di-GMP; the L634P mutation in DgcH could mimic a c-di-GMP bound state to cause the observed reduction in c-di-GMP. We cannot discount the possibility of additional regulatory mechanisms, since numerous missense and in-frame deletion mutations were detected across the coding sequence of DgcH (Fig. 5). As already discussed above, the V459G mutation greatly increased c-di-GMP production in lineage 1 (Fig. 4), and in lineage 27, the Q454P mutation caused a shift to the D phenotype then reverted back to the M phenotype through another missense mutation (Q148P) (Table S1). Similar to DgcH, specific missense mutations in other DGCs caused bi-directional shifts and most activating mutations were found outside their conserved catalytic domain (Fig. 5). We also detected the I-site in other DGCs, with MorA being the lone exception, but no associated mutations were observed (Fig. 5). To the best of our knowledge, no functional studies have been carried out to characterize the bioinformatically predicted domains of these DGCs beyond their catalytic domain. We have clearly established that our mutations impact c-di-GMP production, which should stimulate future studies to characterize their catalytic and/or regulatory consequences.

**Figure 5.**
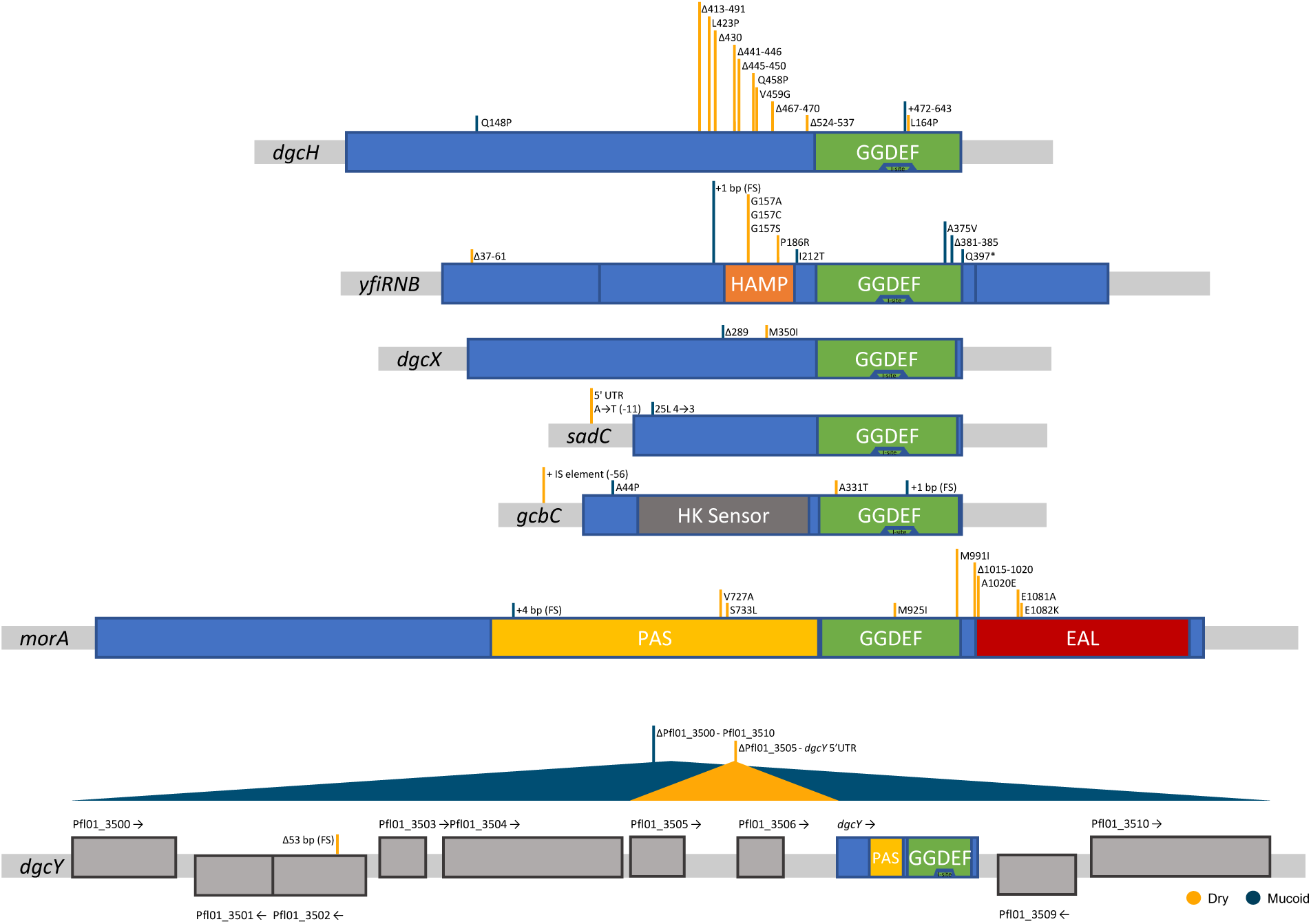
Alignment of our mutation data to annotated domains in known or predicted DGCs. Blue boxes represent the protein sequence and their flanking grey boxes represent untranslated regions. Each image represents a single DGC as indicated, with the exception of *yfiRNB* and *dgcY*. *YfiRNB* shows three blue boxes with each box representing YfiR, YfiN (DGC), or YfiB, respectively. *DgcY* shows a cluster of 10 genes with nine hypothetical genes of undetermined function and one bioinformatically predicted DGC (*dgcY*). Domain data were derived from CDD (NCBI): GGDEF = c-di-GMP catalytic domain, PAS = sensory signaling domain, EAL = phosphodiesterase hydrolysis domain, HAMP = transmembrane relay domain, I-site = c-di-GMP allosteric regulation site (shown inside GGDEF), HK Sensor = histidine kinase sensory domain. Mutations that stimulate c-di-GMP production are indicated by orange horizontal bars and those that reduce c-di-GMP production are indicated by blue horizontal bars.

### Mutations in genes with previously unknown associations with c-di-GMP production integrate with Wsp or other DGC mutations to modulate c-di-GMP levels

We also observed mutations in bioinformatically annotated genes with no known associations to c-di-GMP modulation, but with clear associations with Wsp components and other DGCs. This pattern is best represented in lineage 5 (Fig. 4). Like most lineages, mutations initially accumulate within *wsp* genes and deregulate the Wsp system. Subsequent strains D 5.5 and M 5.6 carry mutations that activate then deactivate the DGC SadC, respectively. Surprisingly, strain 5.7 returns this lineage to the D phenotype through a missense mutation in IlvH (G150S), which is predicted to function in the biosynthesis of branched chain amino acids (BCAA). We observed three independent missense mutations in IlvH across our study that exclusively cause a phenotypic shift from M to D and reduced motility (Table S1). Accordingly, quantification of c-di-GMP shows that each of these three mutations significantly increases c-di-GMP respective to their immediate M parent (Fig. 6). These results indicate that a previously unknown relationship exists between BCAA biosynthesis and c-di-GMP production, and we explore this relationship in detail below.

**Figure 6.**
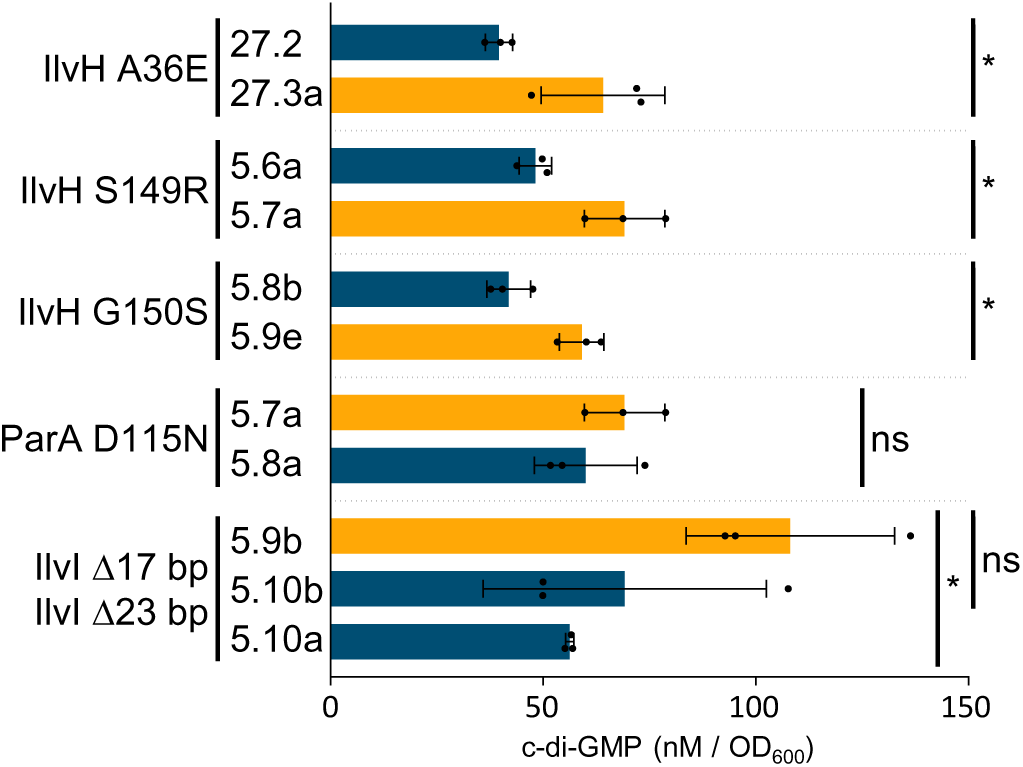
C-di-GMP levels fluctuate with BCAA mutations. C-di-GMP measurements in triplicate mean ± SD. Raw c-di-GMP values of each mutant and its respective parent are reported in Table S1. Causal mutation of each phenotypic shift is indicated on the left. D strains are shown in orange and M strains are shown in blue. (Student’s t-test: ns = non-significant, * = p < 0.05).

Lineage 5 returns to the M phenotype in strain 5.8 through a missense mutation in the hypothetical protein Pfl01_0730 (D115N) (Fig. 4). This protein, hereon ParA (<u>P</u>AP2 <u>a</u>ssociated c-di-GMP regulator), shares homology with the PAP2 superfamily of proteins, suggesting that it may be involved in phospholipid turnover and impact the bioavailability of phosphate within the cell (36). Although we found the relative decrease in c-di-GMP in strain 5.8 to be not statistically significant from its parent likely due to technical noise (Fig. 6), the observed mutation clearly drives the phenotypic transition from D to M. Furthermore, if this mutation does not impact c-di-GMP levels, then we would not expect a further transition through elevated c-di-GMP production. Indeed, we captured two independent transitions back to the D phenotype, both with substantially increased c-di-GMP levels (Fig. 4). Strains 5.9a and 5.9b are sister strains that have developed unique chimeric Wsp proteins. Both chimeric proteins (WspA::D and WspB::D) contain identical partial segments of WspD commencing at residue 158, situated immediately adjacent to the in-frame deletion detected in their common ancestral strain 5.4, which previously deactivated the Wsp system (Fig. 4). To determine whether or not these chimeric proteins alone are sufficient to drive the phenotypic shift from M to D, we replaced the corresponding native Wsp proteins in the ancestral M strain with each chimeric protein. In parallel, we also replaced the corresponding naturally modified Wsp proteins in the immediate parental M strain 5.8 with each chimeric protein. Both engineered strains in the ancestral M background retained the M phenotype, but both engineered strains in the M strain 5.8 background recapitulated the D transition that was observed in strains 5.9a and 5.9b (Table 1). These results indicate that the two chimeric Wsp proteins require additional mutation(s) observed within this lineage to produce c-di-GMP. Both the WspA::D and WspB::D chimeras have lost WspC, which is not surprising since the methylation sites of WspA were removed previously in the D strain 5.3. Similarly, WspB is completely removed in the WspA::D chimera and nearly 75% of WspB has been removed in the WspB::D chimera. At this stage, it remains unclear whether or not the observed mutations in *ilvH* and/or *parA* are required for the function of the two Wsp chimeras. However, we present additional evidence below that strengthens the relationship between IlvH and the Wsp system.

The WspB::D chimera strain 5.9b was next used to evolve M progeny strains. Sister strains 5.10a and 5.10b were isolated and each were found to carry frameshift deletions in *ilvI* that decreased c-di-GMP levels (Fig. 4). The two *ilvI* mutants carry a deletion of 17 or 23 nucleotides that start at the same position, which should abolish IlvI’s function by causing a frameshift. IlvI and IlvH form a complex that catalyzes the first step of BCAA biosynthesis, where IlvH serves as the regulatory subunit and IlvI serves as the catalytic subunit (37). IlvH in *E. coli* has been bioinformatically predicted to respond to c-di-GMP levels due to the presence of the inhibitory I-site (35), which we also detected in most DGCs (Fig. 5). We identified that IlvH of *P. fluorescens* Pf0-1 also contains the I-site RXXD motif at residues 103-106. None of the independently identified IlvH mutations (A36E, S149R, and G150S) in our study fall within this motif, but all three mutations significantly increase c-di-GMP levels (Fig. 6). Beyond the potential allosteric regulation of IlvH by c-di-GMP, there is no known relationship between BCAA biosynthesis and c-di-GMP production to the best of our knowledge. However, we suspected that the IlvH mutations, including the G150S mutation observed earlier in this lineage, increase BCAA production to stimulate c-di-GMP production through an unknown mechanism, since we subsequently observed frameshift mutations in the catalytic subunit (IlvI) that returns this lineage back to the M phenotype with reduced c-di-GMP levels.

The frameshift mutations in *ilvI* should lead to BCAA auxotrophy, which is likely masked by the nutrient-rich medium utilized in our experimental evolution system. To test this prediction, we grew strains 5.10a and 5.10b in a chemically defined minimal medium without BCAA. We did not observe any growth unless BCAA was exogenously supplemented (Fig. S3), confirming that our *ilvI* mutants are unable to synthesize BCAA. We next tested if exogenous supplementation of BCAA alone could restore c-di-GMP production in the *ilvI* mutant (M strain 5.10a). Since the *ilvI* mutant cannot grow in the minimal medium without BCAA supplementation, we also included our prototype M and D ancestors (Fig. 1A) as relative controls in addition to the *ilvH* mutant (D strain 5.7) and the *parA* mutant (M strain 5.8) from the same lineage. Exogenous supplementation of BCAA in the *ilvI* mutant indeed restored c-di-GMP production at comparable levels to both D and the *ilvH* mutant (D strain 5.7) (Fig. 7). In contrast, c-di-GMP levels remained the same in both M and the *parA* mutant (M strain 5.8) regardless of BCAA supplementation (Fig. 7). These results show that overabundance of BCAA does not correlate with c-di-GMP production and defeats our initial prediction that the missense mutations in *ilvH* increase BCAA production to stimulate c-di-GMP production. Although BCAA supplementation restores c-di-GMP production in the *ilvI* mutant strain in the minimal medium, the reduction of c-di-GMP levels observed in the same *ilvI* mutant under our nutrient-rich experimental evolution conditions (Fig. 6) is not likely associated with losing the ability to synthesize BCAA. Accordingly, reconstructing the two frameshift *ilvI* mutations independently in the ancestral D strain did not cause a shift to the M phenotype, but the initially observed shift to the M phenotype was recapitulated when these mutations were introduced into their direct parental D strains (Table 1). Since IlvI forms a complex with IlvH, the observed frameshift mutations in *ilvI* likely decouple the complex, and consequently impact IlvH’s function to modulate c-di-GMP production. Lineage five returns to the D phenotype in four sister branches following the *ilvI* mutations through mutations in *wspE* or *dcgH* (Fig. 4). As already discussed above, we commonly observed activating *wspE* mutations when the upstream signaling Wsp components are disrupted, and the in-frame deletion in *dgcH* is also identical to the deletion reported in the D strain 4.1. These four independent transition steps into the D phenotype clearly indicate that the catalytic activity of IlvI is not required for c-di-GMP production.

**Figure 7.**
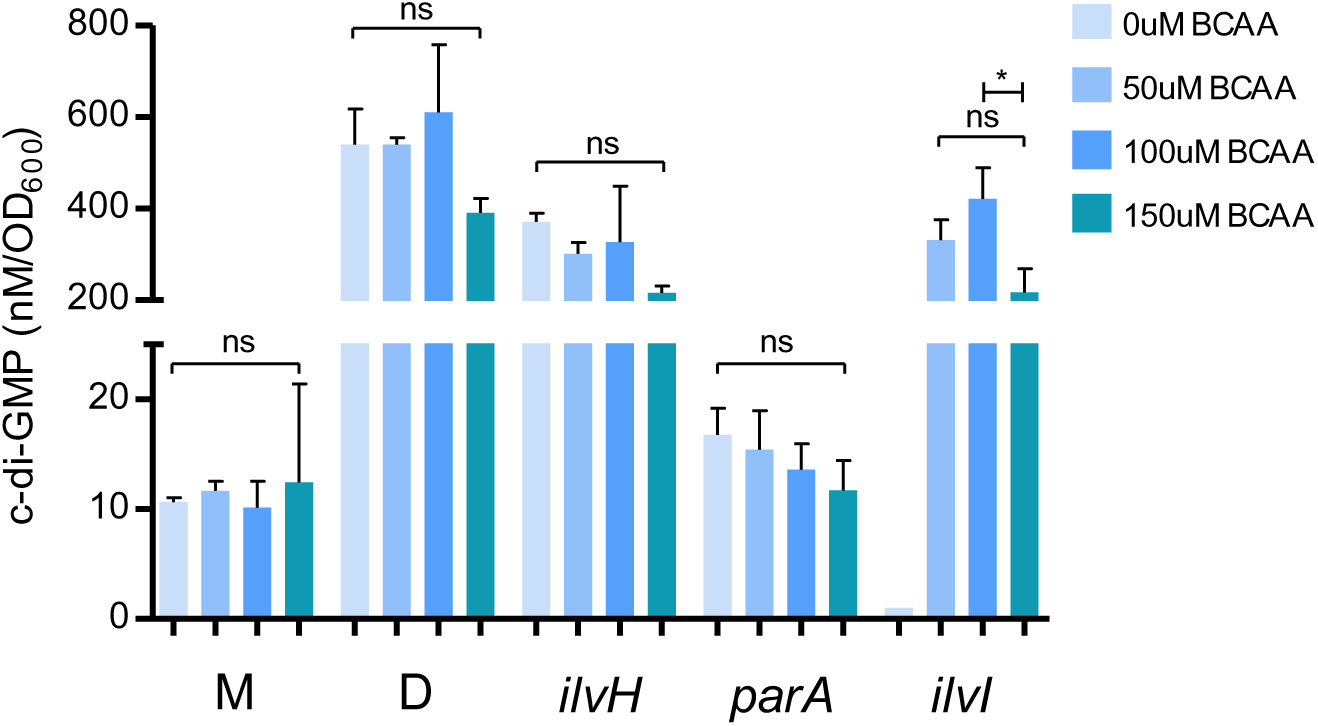
BCAA supplementation stimulates c-di-GMP production in a BCAA auxotroph with the M phenotype, but does not impact c-di-GMP production in non-BCAA auxotrophs with the same M phenotype. Strains were grown in the chemically defined PMM medium with 0, 50, 100, or 150 uM BCAA supplementation (valine, leucine, isoleucine). C-di-GMP was extracted and quantified via LC-MS/MS in triplicates and plotted as the mean with STD (ANOVA; Tukey’s HSD Post Test: ns = non-significant, * = p < 0.05). M and D refer to the prototypical M and D strains in Fig. 1A. The *ilvH* mutant (D strain 5.7) produces c-di-GMP levels similar to the D strain control and the *parA* mutant (M strain 5.8) produces c-di-GMP levels similar to the M strain control. These relative patterns are consistent with the c-di-GMP quantification data reported in Fig 6 under nutrient-rich conditions. Exogenous supplementation of BCAA across all concentrations does not significantly impact c-di-GMP production. There is no data for the *ilvI* mutant at 0 uM BCAA, since this strain fails to grow in PMM unless BCAA is exogenously provided. However, the *ilvI* mutant (M strain 5.10a) produces c-di-GMP levels comparable to both D and the *ilvH* mutant (D strain 5.7) with BCAA supplementation.

Lineage 27 was commenced with an engineered *ΔwspCD* M strain (Table S1, strain 27.0), where we also observed the shift to the D phenotype through a missense mutation in IlvH (A36E) by restoring c-di-GMP production (Fig. 6). In contrast to the M-shifting ParA (D115N) mutation that followed the D-shifting IlvH (G150S) mutation in lineage 5, the transition from IlvH (A36E) to the M phenotype was associated with a mutation which converts the stop codon of *wspB* with a cysteine residue to extend its open reading frame, but out of frame with *wspE* (both *wspC* and *wspD* were already deleted). A common trend exists between lineages 5 (Fig. 4) and 27 (Table S1) in that multiple mutations first occur in Wsp proteins, which is proceeded by mutations in a non-Wsp DGC (SadC or DgcH, respectively), then unique missense mutations occur in IlvH to stimulate c-di-GMP production. Reconstructing the three IlvH missense mutations in the ancestral M strain does not lead to a shift to the D phenotype, but reconstructing these mutations in their direct parental M strain recapitulates the initially observed shift to the D phenotype (Table 1). The common targets of mutations between the two lineages that converge on IlvH mutations are Wsp proteins, which implies that the c-di-GMP stimulating mutations in IlvH specifically associate with a de-regulated Wsp system. Furthermore, the undoing of the IlvH (A36E)-induced c-di-GMP production by the stop codon mutation in *wspB* implies potential IlvH-Wsp interactions that are unlikely to be purely metabolic in nature.

### A macro assessment of sequential mutation patterns across all 33 parallel lineages captures potential associations within the c-di-GMP regulome

A long-standing question in the field is why microbes possess so many independent DGC and/or PDE machineries. A systematic analysis of genes that are bioinformatically predicted to be DGCs in *P. fluorescens* demonstrated that most have little impact on the intracellular c-di-GMP pool (9), and a significant number of DGCs/PDEs in *E. coli* are present in an inactive form (8). In addition, independent DGCs could regulate a sub-cellularly localized role (38) that could also contribute to the total intracellular pool to regulate diverse physiological pathways across a gradient (39). Our study indicates that many known or predicted DGCs could be independently stimulated to trigger a common phenotypic shift, which appears to operate across a broad range of c-di-GMP levels (Fig. 1C). We also captured mutations in genes with unknown associations with c-di-GMP modulation that significantly impact the c-di-GMP pool (Table S1). Repetitive patterns of sequential mutations across independently evolving lineages could reveal previously unknown connectivity between discrete regulatory and catalytic elements.

To visualize conserved patterns of mutations across our complex dataset, we compiled all sequences of mutation events across the 33 parallel lineages into a single node-edge plot (40) (Fig. 8A). The nodes represent the mutated genes in our experiment. The edges (i.e. lines) of the plot report if the mutation causes an M (blue) or D (orange) phenotype, and the thickness of the edge reflects the frequency of the observed pattern (Table S2). Proteins that are integrated into this plot – meaning that many edges connect to a specific node – collectively form the main ring. Nodes associated with a single edge fall out of this ring, and mutations in these proteins typically occur at the end of a lineage or a branched sub-lineage that was not sequenced. Since we did not sequence any progeny strains from these ten mutants that fall outside of the main ring, we will not discuss them further.

**Figure 8.**
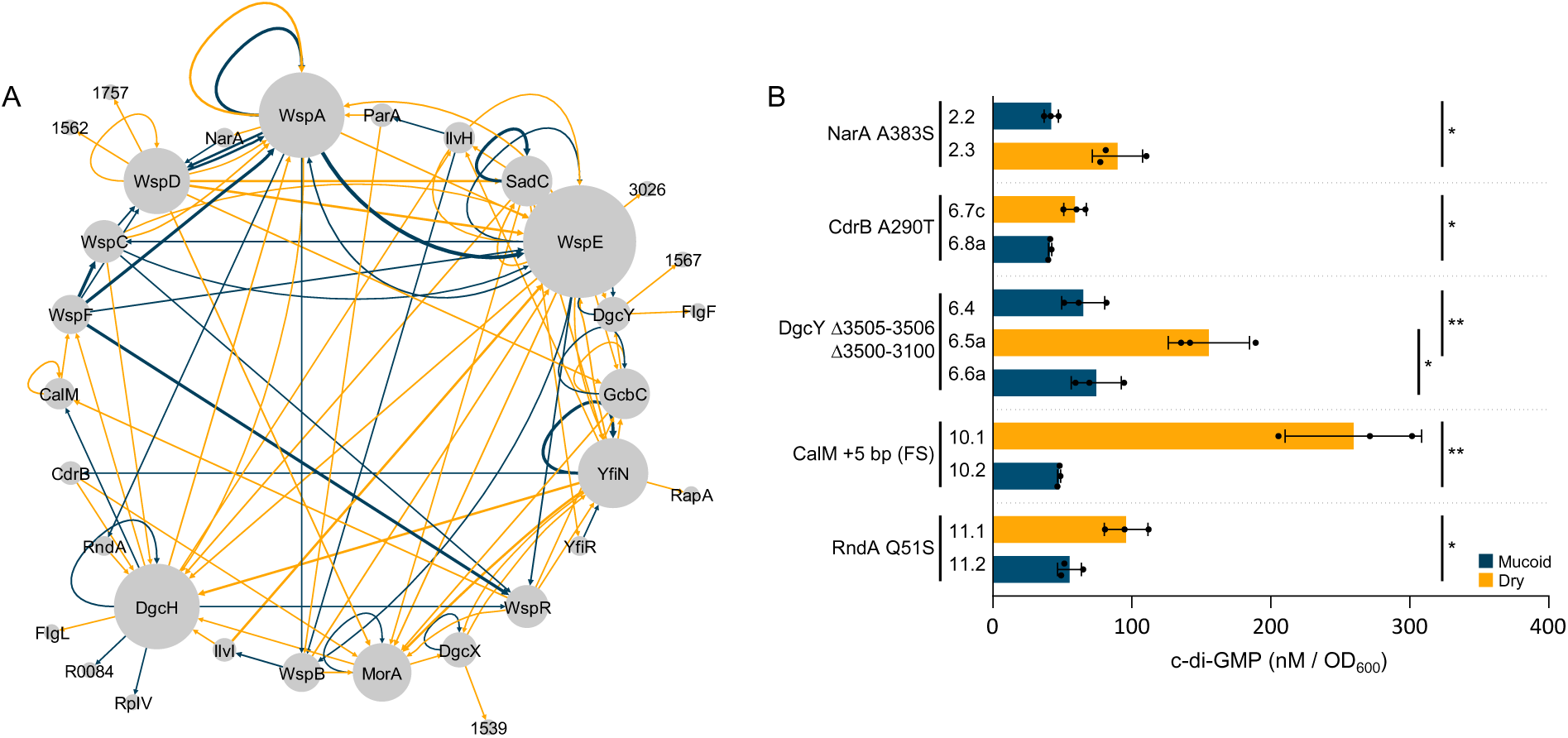
A node-edge representation of 147 unique mutations across 33 lineages and c-di-GMP quantification data of strains with mutations in proteins with significant network connectivity that are not known to be associated with c-di-GMP modulation. A) A node-edge plot reveals focal proteins under selection. Each node represents a protein and each edge represents a relationship where one mutation precedes or proceeds another. Orange edges represent M to D shifts and blue edges represent D to M shifts. The relative size of a node and thickness of an edge is proportional to frequencies within the dataset. Proteins that collectively form the ring represent commonly mutated targets in this study and likely have a substantial role in c-di-GMP regulation. Proteins with limited connectivity fall outside the main ring. B) LC-MS/MS validates that proteins with significant network connectivity, but not previously associated with c-di-GMP, have a significant impact on intracellular c-di-GMP levels. C-di-GMP measurements are reported as the triplicate mean ± SD. C-di-GMP values of each mutant and its respective parent are summarized in Table S1 with the corresponding ID tag. D strains are shown in orange and M strains are shown in blue. (Student’s t-test: * = p < 0.05, ** = p < 0.01).

The largest nodes in Fig. 8A are represented by Wsp proteins, DgcH, and other known DGCs like YfiN, GcbC, MorA, and SadC. The majority of DGCs have blue recursive edges that cause an M phenotype, which indicates that mutations occur in the same protein to shut down its activity, immediately following a mutation that initially stimulated its activity. The high frequency of blue recursive edges associated with each DGC and yellow edges between distinct DGCs collectively indicate that each DGC simply replaces another to restore c-di-GMP production. Blue edges between DGCs are extremely rare, which suggests that individual DGCs do not directly interact with another. Notably, there are several proteins with both yellow and blue edges that are integrated in this plot and have not been shown to have a role in c-di-GMP regulation or production. We next infer potential function of these proteins with respect to c-di-GMP modulation.

#### NarA

(<u>N</u>-ethylammeline chlorohydrolase <u>a</u>ssociated c-di-GMP regulator): Pfl01_3147 is annotated as a hypothetical protein, but we found homology to an N-ethylammeline chlorohydrolase. In strain 2.3, we observe that the A383S mutation in NarA increases c-di-GMP levels (Fig. 8B), which confirms that this hypothetical protein is indeed biologically active. This NarA mutation follows the M-shifted W90L mutation in WspD, which suggests that NarA is either able to restore Wsp signaling or activate other DGCs to increase c-di-GMP levels. We have no reason to suspect that NarA directly catalyzes c-di-GMP since it lacks any c-di-GMP associated domains or motifs. The progeny of this NarA mutant carries a nonsense mutation in *wspD*, reverting this strain back to an M phenotype (Table S1). Thus, the A383S mutation in NarA associates specifically with *wspD* mutations to both restore and shut down c-di-GMP production. NarA is predicted to catalyze N-ethylammeline, a derivative of ammeline, and a recent study theorized that ammeline is likely an essential building block required for the synthesis of cyclic complexes (41). Although the metabolic connections are unknown, there is a strong indication that NarA interacts with WspD at some capacity to modulate c-di-GMP levels.

#### CdrB

(Two-partner secretion system transporter): Pfl01_0652 encodes a protein that shares homology with the transporter protein CdrB in *P. aeruginosa* (PA4624). In *P. aeruginosa*, CdrA and CdrB are encoded in a single operon and form a two-partner secretion (TPS) system associated with biofilm formation (42). CdrA (c-di-GMP regulated TPS A) is an extracellular adhesin known to respond to modifications made by the periplasmic protease LapG which is activated by the transmembrane c-di-GMP receptor LapD (43). It has been reported that high intracellular levels of c-di-GMP enable CdrA to bind and crosslink secreted Psl, thereby increasing cell-cell adhesion (42). *cdrA/B* expression also increases proportionally with intracellular levels of c-di-GMP during biofilm formation (44). CdrB is a transporter with domains that indicate it is a type V secretion system transporter. CdrB is likely involved in the transport of the adhesin CdrA but has no identified role in c-di-GMP modulation. In our lineage 6, the missense mutation YfiN (G157C) increases c-di-GMP production, which is immediately followed by the CdrB A290T missense mutation to decrease c-di-GMP levels (Fig. 8B). This suggests that an unaltered CdrB or its native transport activity is necessary for YfiN’s DGC activity. Mutations that follow the CdrB mutation occur in *morA* or *dgcH*, each increasing c-di-GMP as indicated by the D-phenotype (Table S1). Therefore, the negative impact of our CdrB mutation on YfiN could be recovered through the genetic activation of other DGCs, which could also reflect interactions with CdrB.

#### CalM

(<u>c</u>-di-GMP <u>a</u>ssociated <u>L</u>ys<u>M</u>): The hypothetical gene Pfl01_1895, hereon *calM* (<u>c</u>-di-GMP <u>a</u>ssociated <u>L</u>ys<u>M</u>) is a commonly mutated gene in our study (Table S1). The encoded protein contains domains with homology to the LysM peptidoglycan-binding superfamily and contains a FimV C-terminal transmembrane domain that is implicated in modulating motility (45, 46). The LysM proteins are identified by the lysin motif which is associated with cell lysis. The presence of the LysM and FimV domains suggests that this protein may respond to changes in peptidoglycan stress, which is essential for maintaining cell shape and integrity. C-di-GMP is a well-known regulator of cellular morphology in *Vibrio cholera* and *Caulobacter crescentus* (13, 47). In addition, surface-sensing through the Wsp system in *P. aeruginosa* was recently demonstrated to be associated with cell envelope stress (28). We observed a duplication of five nucleotides within *calM* that results in a significant decrease in c-di-GMP (Fig. 8B). Two repetitive clusters of CAGCG shifts to three clusters to cause a frameshift. This mutation follows a *dcgH* in-frame deletion that dramatically increases c-di-GMP production, indicating that CalM is necessary for DgcH’s activity. Remarkably, three independently evolved progeny strains in this lineage carry mutations that remove the initial five nucleotide insertion, returning each strain back to the D phenotype likely by restoring DgcH’s activity (Table S1).

#### RndA

(RND transporter): Pfl01_2749 is bioinformatically predicted to encode an RND transporter. RND transporters function as efflux pumps to remove toxins, antibiotics, quorum sensing (QS) molecules, and other metabolites (48). We observed that the nonsense mutation Q51* in Pfl01_2749, hereon *rndA*, significantly decreases c-di-GMP levels (strain 11.2, Fig. 8B). A recent study speculates that QS molecules contribute to a feedback loop to inhibit c-di-GMP production (49). Thus, an active RndA could potentially modulate this feedback loop by expelling certain effector compounds to stimulate c-di-GMP production. The *rndA* mutant shifts to the D phenotype through an in-frame deletion that removes amino acid residues 524-537 in DgcH. This specific mutation was observed in multiple lineages to exclusively stimulate c-di-GMP production.

#### DgcX and DgcY

(Bioinformatically predicted DGCs): Pfl01_3550 and Pfl01_3508 are both bioinformatically predicted to encode DGCs, but their enzymatic function has not yet been demonstrated. Here, we establish that they are both functionally active in c-di-GMP catalysis and annotate them as *dgcX* and *dgcY*, respectively, since genes *dgcA* through *dgcW* already exist. Pfl01_3550, hereon *dgcX*, is a commonly mutated gene in our dataset which increases c-di-GMP (Table S1). We observed a blue recursive loop in the node-edge plot for DgcX, which is consistent with other known DGCs (Fig. 8A). We isolated two independently evolved D mutants with the same M350I mutation, and an M mutant with an in-frame deletion of L289. Phenotypic and motility data support that the observed mutations in *dgcX* influence the intracellular c-di-GMP pool. The genomic location of *dgcY* is in a region of hypothetical genes that appear to form an operon. Interestingly, our *dgcY* mutant strains carry large deletions of the hypothetical genes within this genomic region (Fig. 5). The deletion of Pfl01_3505, Pfl01_3506, and the 5’ UTR region of *dgcY* results in a significant increase in c-di-GMP (Strain 6.5a, Fig. 8B), which likely replaces the native promoter region of *dgcY* with that of Pfl01_3505. C-di-GMP is significantly decreased in the progeny strain 6.6a (Fig. 8B) after the deletion of Pfl01_3500-Pfl01_3510, which includes *dgcY*. It is unclear whether or not the encoded proteins of the hypothetical genes that flank *dgcY* are functionally related to DgcY. We had initially suspected that these hypothetical genes could encode a complex signal transduction system like Wsp, but we did not detect any homologous Wsp domains or other c-di-GMP associated motifs.

To confirm whether the observed mutations in these atypical c-di-GMP associated genes alone could modulate c-di-GMP levels, we reconstructed representative mutations in their respective ancestral backgrounds. None of the engineered strains produced a phenotypic shift, but reconstructing the same mutations in their direct parental strains recapitulated the initially observed phenotypic shifts (Table 1). These results clearly indicate that specific mutations in the associated DGCs are first required to impact c-di-GMP levels. Although functional studies are required to prescribe specific mechanisms, diverse cellular processes appear to be integrated with numerous catalytic machineries to modulate the c-di-GMP pool in *P. fluorescens*, and the missense and in-frame deletion mutations observed in Wsp proteins and other DGCs (Fig. 5) could reflect potential interaction sites.

## Conclusion

Although we have extensive knowledge on how specific enzymes make or break c-di-GMP and how this molecule interacts with a large number of regulatory elements to modulate diverse physiological processes, relatively little is known about how C-di-GMP production itself is regulated. This study takes advantage of a repeatedly evolving social trait in *P. fluorescens* to demonstrate how numerous proteins could independently and collectively produce remarkable sequences of genetic innovation to oscillate c-di-GMP levels. Our extensive library of functionally critical residues reinforces the current model of the Wsp system, which heavily draws functional analogies to the *E. coli* chemotaxis system and particularly lacks domain resolution in WspB and WspD. Beyond the Wsp system, we have confirmed that several bioinformatically predicted DGCs are indeed enzymatically active and identified unique missense and in-frame deletion mutations in multiple DGCs that fall outside their conserved catalytic domain. These mutations likely reflect regulatory domains that are critical for modulating enzymatic activity. We have also uncovered a set of unexpected or hypothetical proteins that clearly have the capacity to influence c-di-GMP turnover. Three missense mutations in *ilvH* were identified to independently stimulate c-di-GMP production, which appears to be independent of IlvH’s primary role in BCAA biosynthesis, but rather operates through an unknown interaction with the Wsp system. We have identified the same I-site in IlvH as in most DGCs, which suggests that IlvH physically interacts with c-di-GMP. Similarly, CdrB and RndA homologs in other organisms have been implicated to respond to c-di-GMP levels, but they have never been associated with modulating c-di-GMP production. We have confirmed that the mutations observed in these proteins alone do not impact c-di-GMP levels, but do so specifically in the presence of genetically modified DGCS or the Wsp signal transduction system.

An emerging concept here is that the intracellular pool of c-di-GMP, whether it be subcellularly localized or in sum, appears to be modulated through the catalytic activities of many independent DGCs that are in tune with diverse proteins. Such intricacies are masked by the activities of DGCs with high output, but they are clearly hardwired in the genome of *P. fluorescens* and reflect a complex and underappreciated interconnectivity. Similar relationships could also manifest in other *Pseudomonas* and related species that share the same DGCs and unexpected proteins described in this study. In particular, very little is known about the functional domains of most DGCs beyond their conserved catalytic domain. The vast majority of the missense mutations or in-frame deletions captured in this study occur outside of the catalytic domain to clearly activate or shut down c-di-GMP production. Our extensive library of functionally important mutations presented here should facilitate mechanistic studies of phylogenetically conserved DGCs that remain heavily under-explored.

## Material and Methods

### Strains and culture conditions

All *Pseudomonas fluorescens* Pf0-1 (50) derivatives associated with parallel experimental evolution are described in Table S1 and engineered strains are described in Table 1. Strains were routinely grown in Lennox LB (Fisher BioReagents), *Pseudomonas* Minimal Medium (PMM) (51), or on *Pseudomonas* Agar F (PAF) (Difco). *Pseudomonas* F (PF) broth (a non-agar variant of PAF) was prepared with the following: pancreatic digest of casein 10 g/L (Remel), proteose peptone No. 3 10 g/L (Remel), dipotassium phosphate 1.5 g/L (Sigma-Aldrich), and magnesium sulfate 1.5 g/L (Sigma-Aldrich). Cultures were incubated at 30°C and shaken at 250 rpm when applicable.

### Parallel experimental evolution

Pure cultures of the ancestral M strain or engineered M strains were grown overnight in 5mL LB broth and 25uL volumes were spotted on PAF plates, then incubated at room temperature for 3 to 8 days to allow spreading fans to emerge. To ensure consistency across experiments, PAF plates were always prepared and left to dry at room temperature for 2 days before being inoculated. One fan was randomly sampled from each colony and streaked out on LB plates in duplicate. Sampling a fan always produced two strains with either a mucoid (M) or dry (D) colony phenotype. A single isolated colony exhibiting the D phenotype was randomly selected to start a new liquid culture and also frozen down. This process was repeated to sample six independent fans (i.e. six lineages). Three of the six independently isolated D strains were randomly selected and cultured as above to start the next round of experimental evolution. We repeated the same sampling process as above except that new M phenotypes were isolated instead. We bidirectionally selected M and D phenotypes for as many as 13 rounds for a single lineage, but the number of sampled fans at each step was reduced over time in a varying manner across independent lineages. In total, approximately 600 independently evolved isolates were frozen down across 33 parallel lineages. Every isolate was subjected to a motility assay in an LB plate with 0.25% agar, and the prototype M and D ancestors were used as controls (24). In addition, 25uL of pure cultures of each isolate, the ancestral strain with the opposite phenotype, and a 1:1 mixture of both were spotted onto PAF plates in triplicate and assessed for cooperative radial spreading (Fig. 1A) to filter out hyper-motile and non-cooperative mutants (24). Colony diameters were measured and recorded after four days of incubation at room temperature and considered cooperative if the measured diameter of the 1:1 colony exceeded both the diameters of the isolated mutant strain and the respective ancestral strain. Isolates that failed this test were removed from this study.

### Reconstruction of naturally derived mutations in ancestral and direct parental backgrounds

Experimentally evolved isolates were whole genome sequenced to identify the causal mutations as described below. We reconstructed select naturally acquired mutations in the respective ancestral background and also in the direct parental background using previously described methods (24). PCR primers were developed from the sequence data to amplify mutations of interest and are reported in Table S3. Isolates were grown overnight and underwent DNA extractions (Qiagen). Dream Taq DNA Polymerase (ThermoFisher) was used in PCR reactions with purified DNA to generate insert fragments carrying the mutations which ranged in size from 456bp to 934bp. PCR was carried out using the recommended conditions of the Dream Taq amplification kit for amplicons below 2000bp. Inserts were cloned into the pGEM-T Easy vector system (Promega) and then subcloned into the *Not*I site of the suicide plasmid pSR47s (52). Suicide plasmids were heat shocked into *E. coli* S17.1λ*pir*, mated with the target strains reported in Table S3 on LB plates at 30°C for 16 hours, then plated onto PMM plates supplemented with 50 ug/ml kanamycin. We observed that *ilvI* mutants could not grow on PMM and therefore replaced the PMM selection plate with an LB plate supplemented with 50 ug/ml kanamycin and 100 ug/ml ampicillin for *ilvI* mutant constructs. Isolated colonies were grown overnight, serially diluted and plated out on LB plates supplemented with 15% sucrose (w/v) for counterselection. Isolated colonies were placed in cryo-storage as described above. Crude DNA extracts were made for each strain where 100uL of overnight culture was boiled at 100°C, centrifuged at 15,000 RCF for 30 seconds, and 10uL of supernatant was used for PCR using the primers reported in Table S3. PCR products underwent gel extraction and subsequent sanger sequencing (primers in Table S3) to confirm that the mutation was successfully engineered into the target strain. Upon PCR confirmation, pure cultures of cells from cryo-storage underwent DNA extraction (Qiagen) and were submitted for WGS which confirmed that no additional mutations had accumulated in the target strains during mutant construction.

### DNA extraction, whole genome sequencing, and variant calling

Pure cultures were grown overnight in 5mL LB broth at 30°C. 1 mL was spun down to form a pellet for DNA extraction. Pelleted cells were mechanically lysed with the Qiagen TissueLyser LT at 50 oscillations/second for 10 minutes. The Qiagen DNeasy UltraClean Microbial DNA extraction kit was used following the standard operating procedures. Samples were then eluted into 30uL H2O after a 5 minutes incubation at 4°C with the elution media and stored at -20°C prior to shipment. Genomic DNA library preparations were carried out by the Microbial Genome Sequencing Center (Pittsburgh, PA) following standard operating procedures. Pooled and indexed samples were sequenced on the NextSeq 2000 platform and demultiplexed prior to data delivery. The Trimmomatic algorithm was used to group and quality filter paired reads at a Phred score of 20 using a 4bp sliding window. The BreSeq algorithm was used to align the paired reads to the *P. fluorescens* Pf0-1 reference genome (CP000094.2) and evaluate for single nucleotide polymorphisms (SNPs), insertions, deletions, or rearrangements. Output sequence data from BreSeq is compiled as ‘Table S1 – Source Data 1’ and was manually evaluated for each strain to determine the order of mutations for a given lineage.

### Alignment of mutation data to annotated domains

Domain analyses of Wsp proteins and DGCs were conducted and rendered as previously described (17). Peptide sequences of the predicted or known DGCs were obtained from the *Pseudomonas* Genome Database (53). NCBI’s conserved domain database (CDD) was used to evaluate the functional domains (54). Mutation data of Wsp proteins from this study were manually mapped to the rendered consensus sequence of the respective Wsp protein (17).

### C-di-GMP extraction and LC-MS/MS

An isolated colony of each strain was transferred to PF broth and grown overnight. Overnight cultures were diluted to an OD600 of 0.04 in triplicate and incubated for 2-4 hours until the samples reached an OD600 of 0.5, then promptly processed for c-di-GMP extraction. C-di-GMP extraction and quantification procedures followed the protocols established by the Michigan State University Research Technology Support Facility (MSU-RTSF): MSU_MSMC_009 protocol for di-nucleotide extractions and MSU_MSMC_009a for LC-MS/MS, with the following modifications. All the steps of c-di-GMP extraction were carried out at 4°C unless noted otherwise. 1mL of 0.5 OD600 culture was centrifuged at 15,000 RCF for 30 seconds, and cell pellets were resuspended in 100uL of ice cold acetonitrile/methanol/water (40:40:20 (v/v/v)) extraction buffer supplemented with a final concentration of 0.01% formic acid and 25nM c-di-GMP-Fluorinated internal standard (InvivoGen CAT: 1334145-18-4). Pelleted cells were mechanically disturbed with the Qiagen TissueLyser LT at 50 oscillations/second for 2 minutes.

Resuspended slurries were incubated at -20°C for 20 minutes and pelleted at 15,000 RCF for 15 minutes at 4°C. The supernatant was transferred to a pre-chilled tube then supplemented with 4uL of 15% w/v ammonium bicarbonate (Sigma-Aldrich) buffer for stable cryo-storage. Extracts were stored at -80°C for less than two weeks prior to LC-MS/MS analysis at MSU-RTSF. Sample degradation was observed with repeated freeze-thaw cycles, so extracts were never thawed until LC-MS/MS analysis. All samples were evaporated under vacuum (SpeedVac, no heat) and redissolved in 100uL mobile phase (10 mM TBA/15 mM acetic acid in water/methanol, 97:3 v/v, pH 5). LC-MS/MS quantification was carried out with the Waters Xevo TQ-S instrument with the following settings: c-di-GMP-F at *m/z* transition of 693 to 346 with cone voltage of 108 and collision voltage of 33; c-di-GMP at *m/z* transition of 689 to 344 with cone voltage of 83 and collision voltage of 31. C-di-GMP data was normalized to 25nM c-di-GMP-F by MSU-RTSF to account for sample matrix effects and sample loss during preparation. OD600 measurements of the initial samples were used to report the quantified c-di-GMP as nM/OD and visualized with GraphPad Prism9.

### BCAA supplementation assay and c-di-GMP quantification

Pure cultures were grown overnight in 5mL LB broth. 5uL of each culture was transferred into 5mL PMM. Cultures were supplemented with the BCAA valine, leucine, and isoleucine as previously described (55). Assessment of growth during BCAA supplementation contained either 0uM (loading control), 180uM, 200uM, 220uM, 250uM, or 300uM in triplicate. Samples were incubated at 30°C for 24 hours at 250 rpm shaking. Cultures were manually pipetted to resuspend cells and 300uL of culture was transferred to a 96-well black wall clear bottom plate for OD600 quantification. OD was quantified using the SpectraMax plate reader and background (determined from blanks) was removed from each quantified sample. Data analysis and visualization of growth during BCAA supplementation was conducted in Excel. To evaluate the relationship between BCAA and c-di-GMP, pure cultures were grown for 24 hours at 30°C and 250 rpm shaking in PMM liquid media containing either 0uM, 50uM, 100uM or 150uM of supplemented BCAAs. OD600 measurements were taken and samples were diluted to an OD600 of 0.04 into PMM at the same BCAA concentrations. Samples were incubated for 24 hours under the same conditions stated and then sonicated (Time 0.00:20, Pulse 01 01, Amplitude 20%) to disrupt cell aggregates. OD600 measurements occurred every hour until OD600 was approximately equal to 0.25. C-di-GMP analyte extraction was carried out as stated above with the modification that 2mL cultures were used for cell lysis. Quantified values of c-di-GMP are reported as nM/OD600 and were visualized with GraphPad Prism9.

### Node-edge analysis and visualization

Data reported in Table S1 was used to generate the ‘to-from’ table which assesses mutation patterns at the gene/protein level (40). Counts were determined by the frequency of the to-from patterns calculated from Table S1 where a mutation in a given gene/protein (to) occurred after a mutation in another gene/protein (from). This table was uploaded to Cytoscape to visualize the nodes (to genes/proteins) and edges (counts). Default renderings of the plot reported node centrality (commonly mutated targets) through coordinate placement on the graph. This metric was converted to node size with larger nodes representing more commonly mutated targets. Nodes were manually aligned for visual clarity with nodes connected by two or more edges being placed in the main ring and nodes with a single edge being placed outside the ring.

## Acknowledgments

We thank RTSF (Michigan State University) for services and assistance with LC-MS/MS analyses and MiGS (Pittsburgh, PA) for providing genome sequencing. This study was funded by the Charles Henry Leach II Fund 217032 (W.K.) and the Faculty Development Fund G1700030 (W.K.) awarded by Duquesne University.

## Author Contributions

W.K. designed the study, C.K. performed the experiments, C.K and W.K analyzed data and wrote the manuscript.

## Conflict of Interest

The authors declare no conflict of interest.

**Table S1 – Source Data 1. BreSeq summary reports of mutations identified in strains isolated in this study.** Whole genome sequencing data of select strains from the parallel evolution experiments were subjected to variant analysis using the BreSeq algorithm against the published P. fluorescens Pf0-1 genome sequence. Summary files are provided for each analysis where the ‘Strain ID’ reported in Table S1 is represented as the file name.

## Supplementary Information

**Figure S1.**
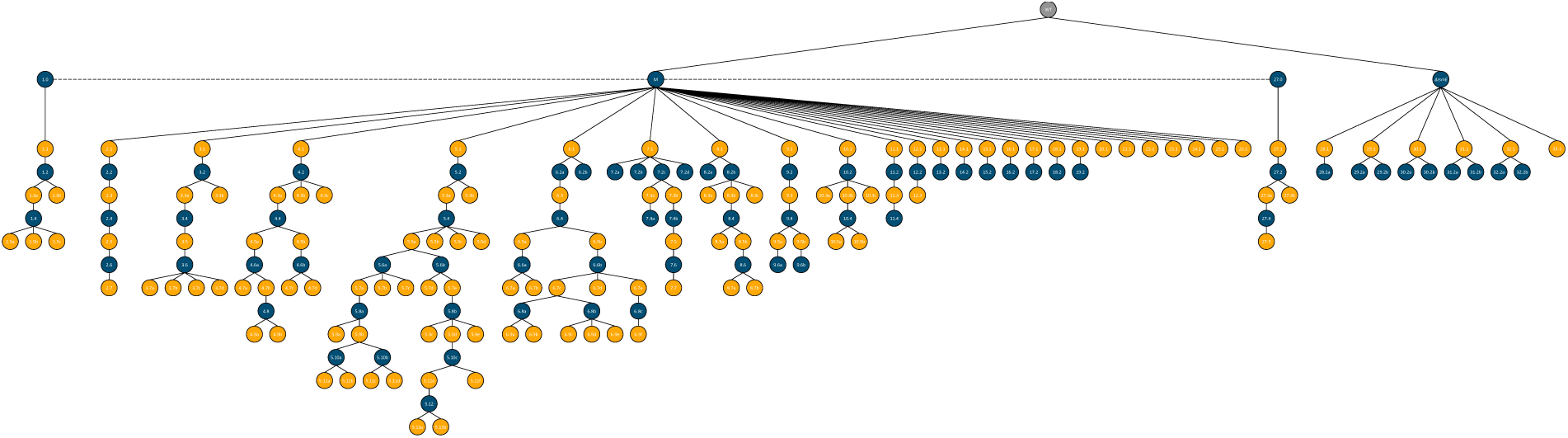
Phylogeny of 191 evolved strains across 33 lineages. Each strain is classified as either D (orange) or M (blue) based on their colony morphology. Corresponding motility data, mutation data, and LC-MS/MS c-di-GMP quantification data are summarized in Table S1.

**Figure S2.**
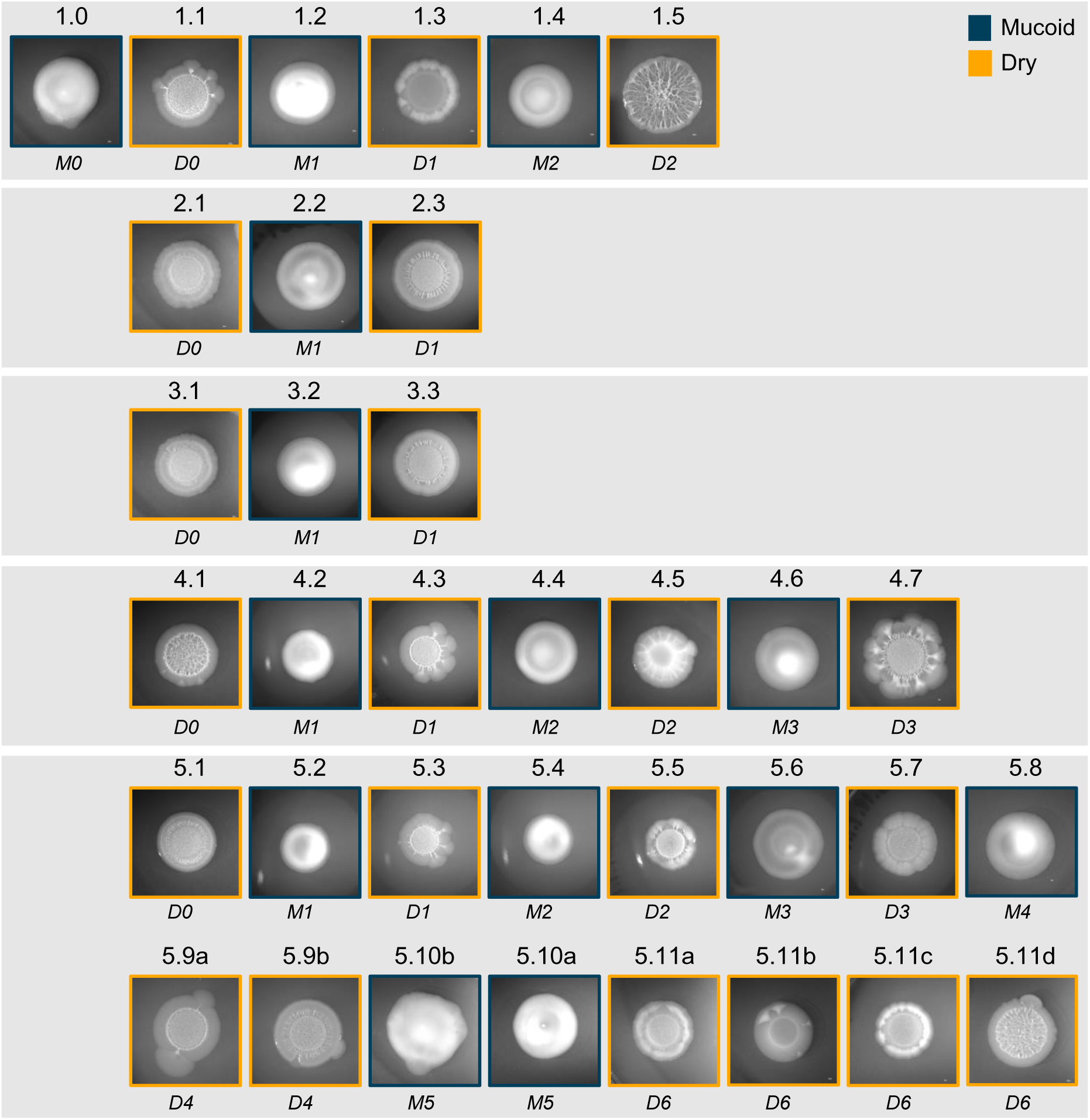
Colony morphologies of evolved strains depicted in Fig. 1B. Each evolved strain is classified either as M (blue) or D (orange) based on their mucoid or dry-wrinkly colony phenotypes, respectively. Motility data and LC-MS/MS c-di-GMP quantification data for each strain are reported in Table S1.

**Figure S3.**
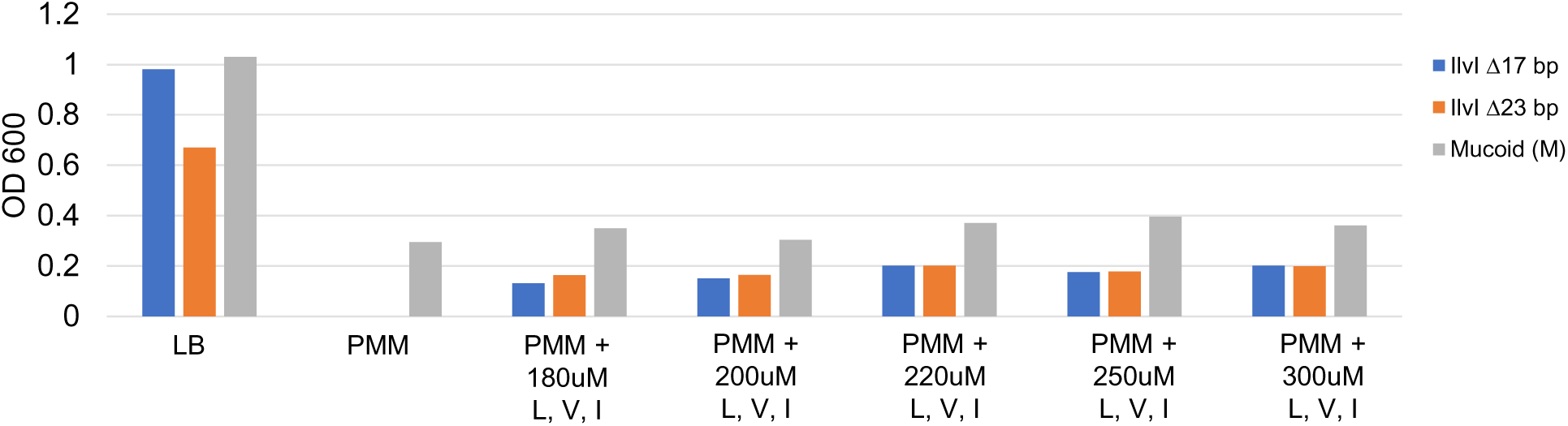
*ilvI* mutants require exogenous supplementation of BCAA. The ancestral M and independently isolated *ilvI* mutant strains were grown in LB (nutrient-rich medium) or PMM (minimal medium) with or without exogenous BCAA supplementation. Both *ilvI* mutants fail to grow in PMM unless supplied with BCAA (leucine (L), valine (V), and isoleucine (I)), and M does not show substantial change in growth with BCAA supplementation.

**Table S1.**
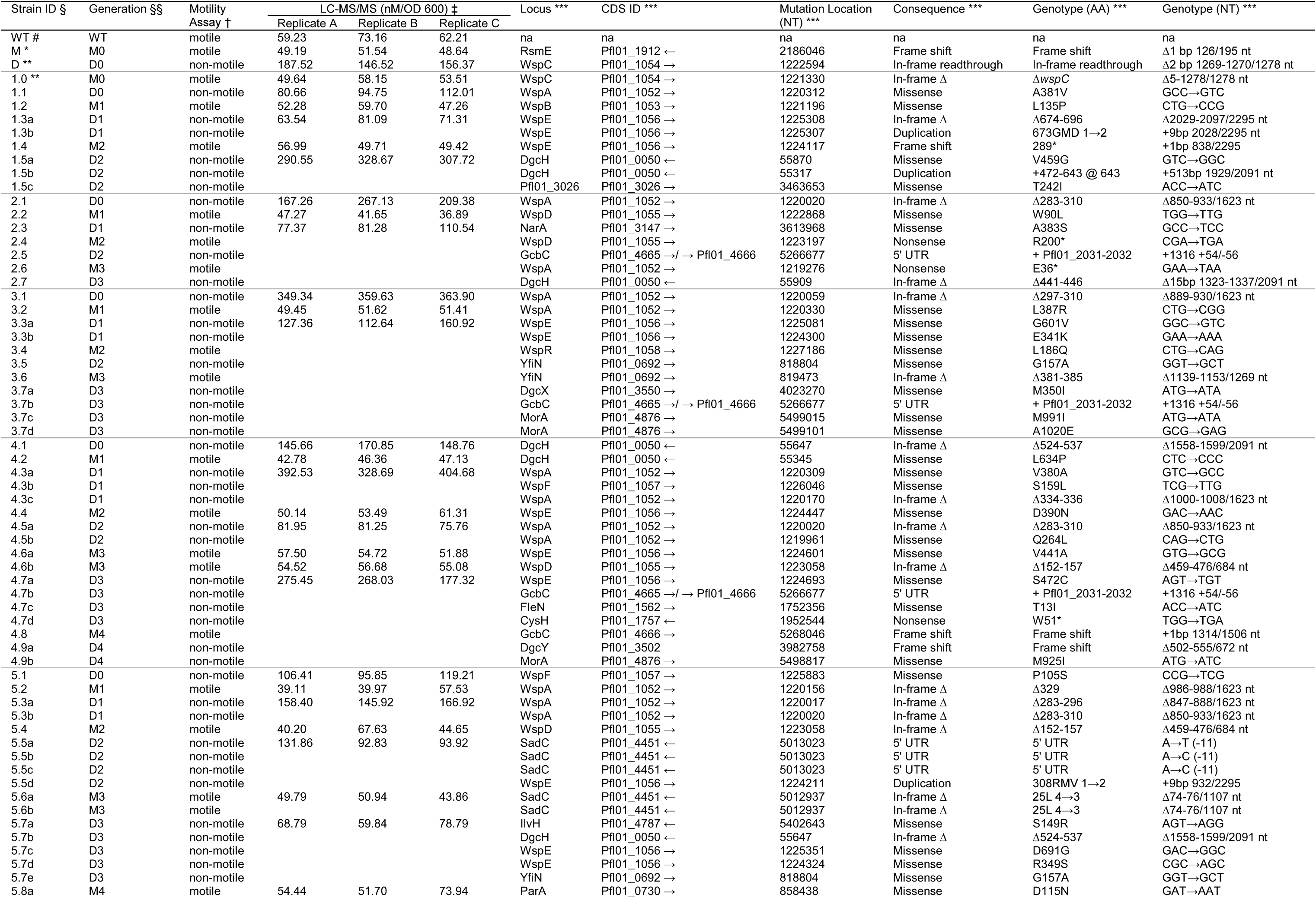
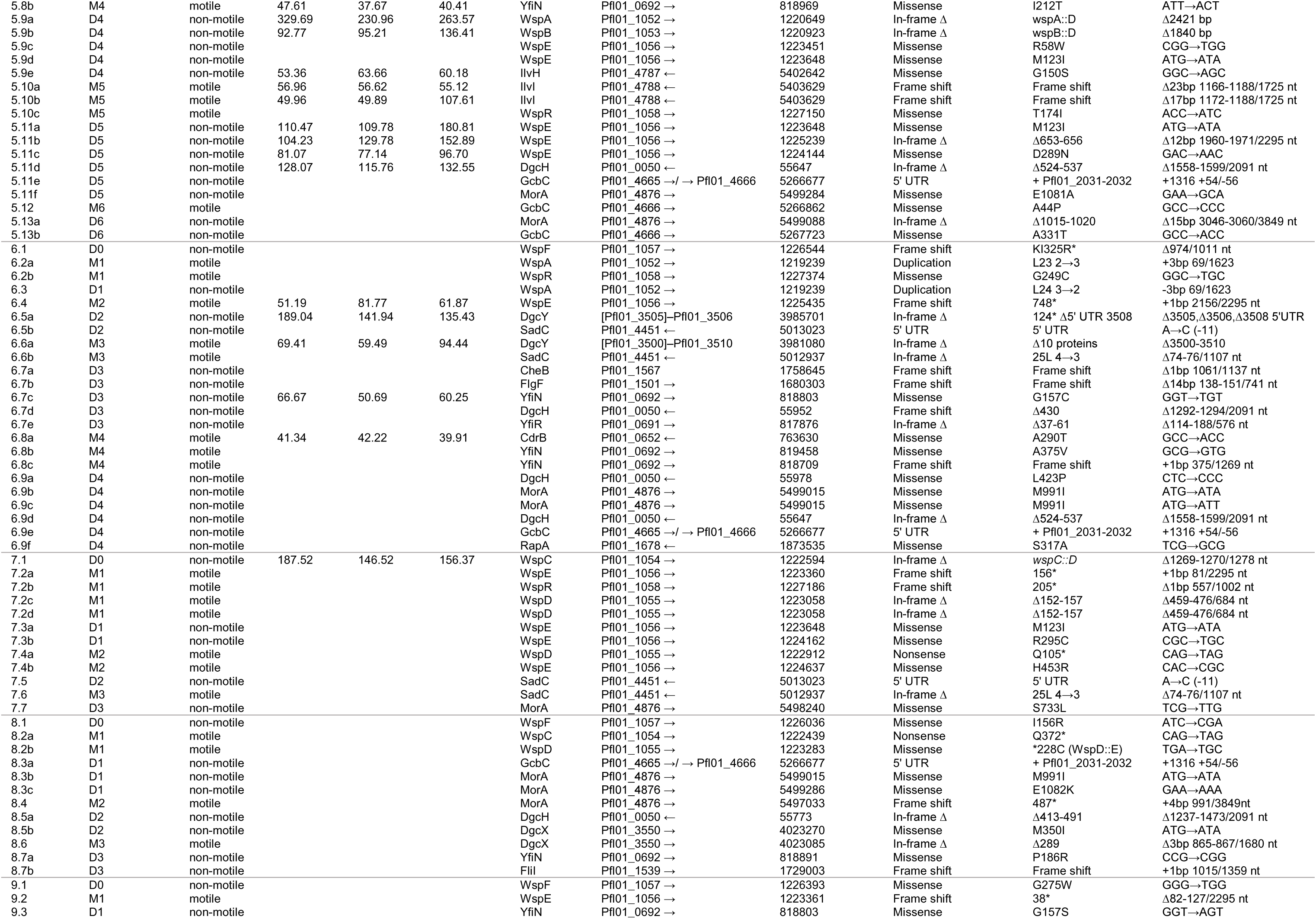
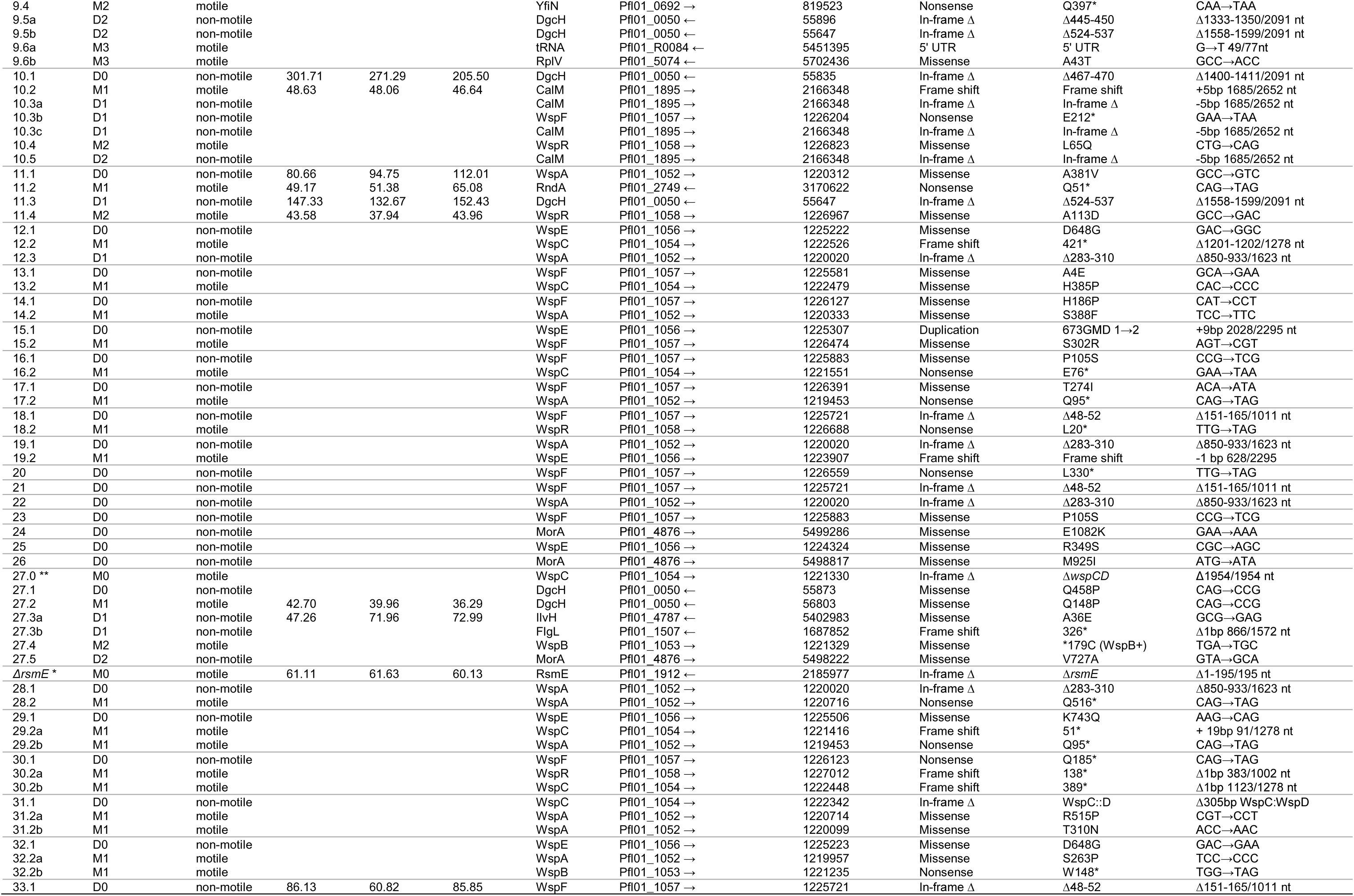
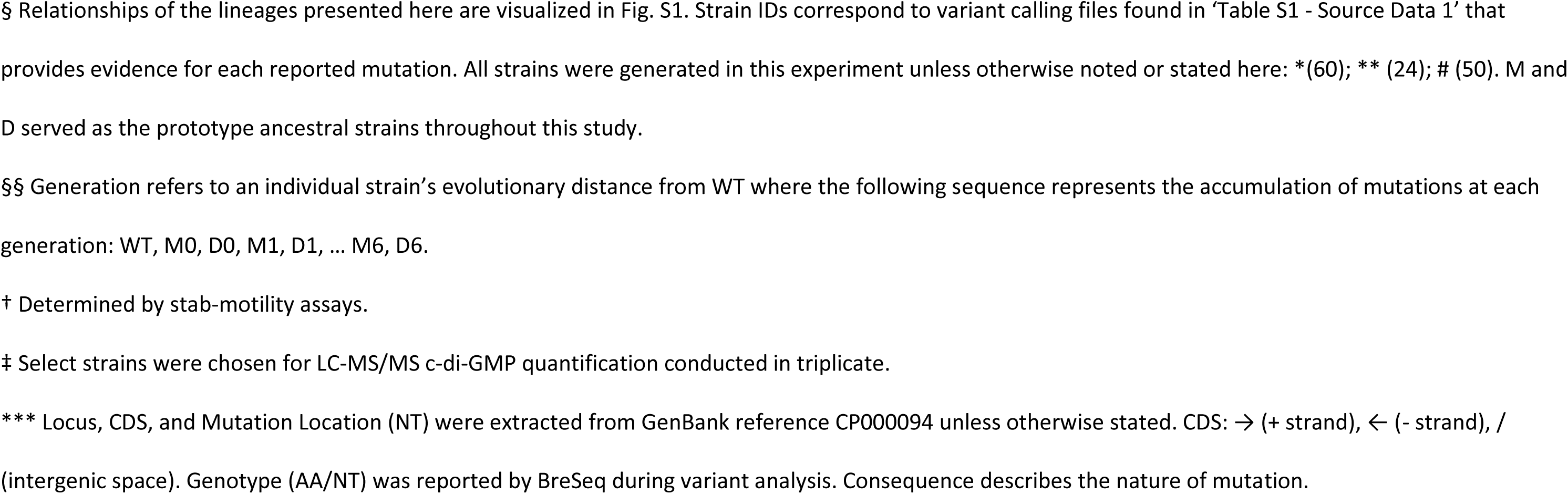
191 strains isolated from the parallel evolution experiment categorized by lineage.

**Table S2.**
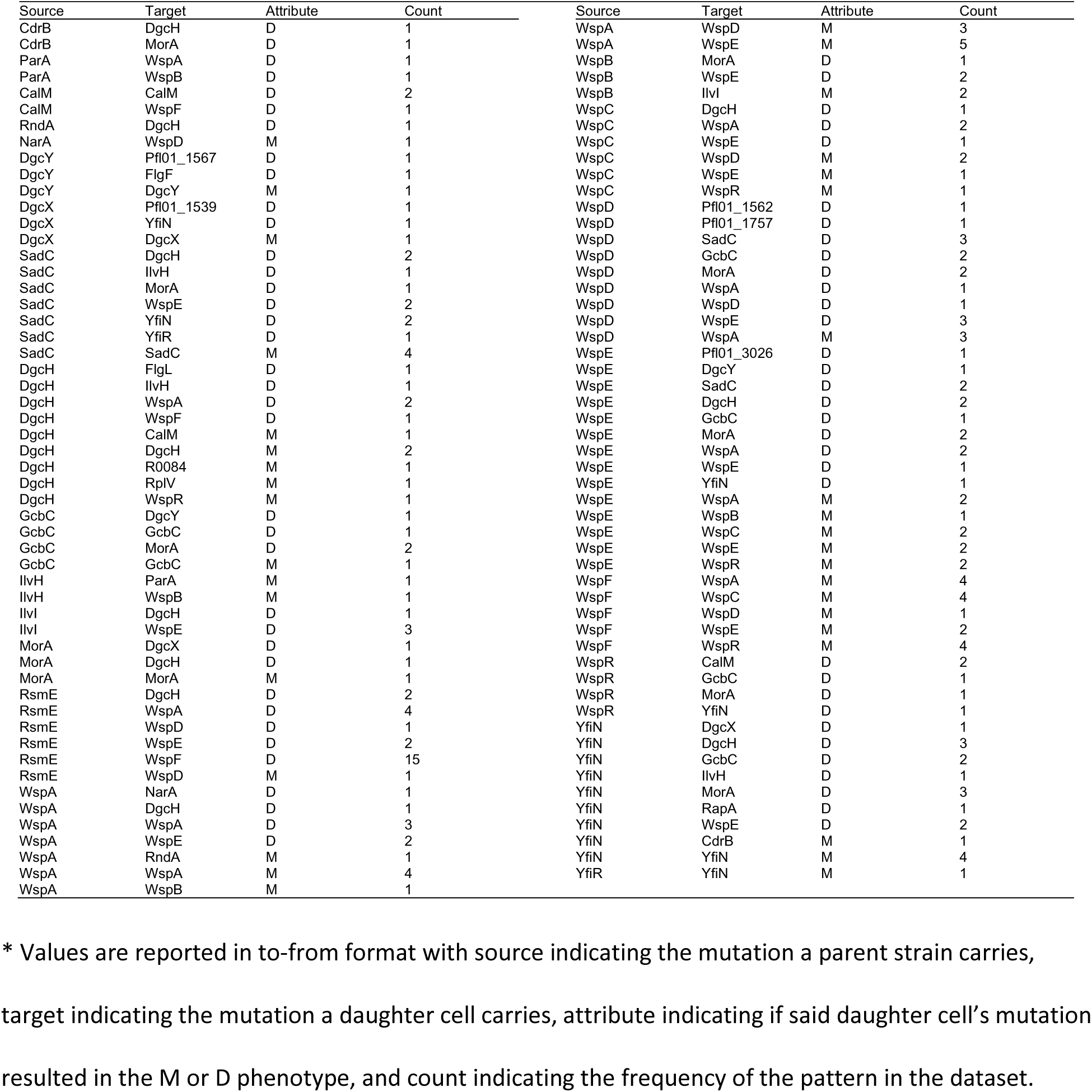
Node-edge plot data matrix.

**Table S3.**
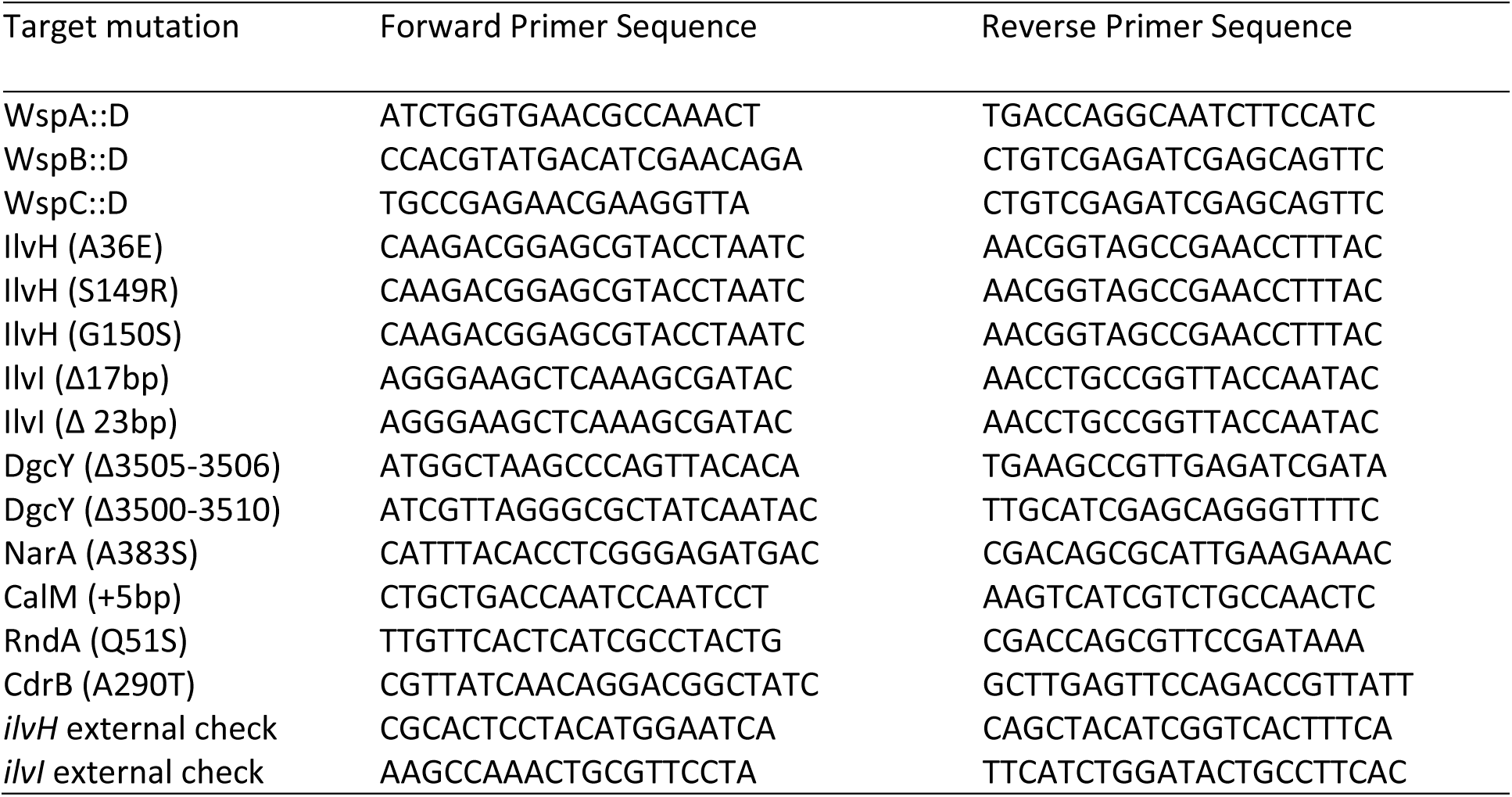
Description of PCR primers used to reconstruct mutations.

